# Cannabinoid Modulation of Central Amygdala Population Dynamics During Threat Investigation

**DOI:** 10.1101/2025.01.21.634174

**Authors:** Farhana Yasmin, Saptarnab Naskar, Luis E. Rosas-Vidal, Sachin Patel

**Author notes:** Corresponding author: Department of Psychiatry and Behavioral Sciences Feinberg School of Medicine Northwestern University 320 E Superior St, 4-490 Chicago, Illinois 60611.

## Abstract

Cannabinoids modulate innate avoidance, threat-reactivity, and stress adaptations via modulation amygdala-associated circuits; however, the mechanisms by which cannabinoids modulate amygdala representation of threat-related behavior are not known. We show that cannabinoid administration increases the activity of central amygdala (CeA) somatostatin neurons (SOM) and alters basal network dynamics in a manner supporting generation of antagonistic sub-ensembles within the SOM population. Moreover, diverging neuronal population trajectory dynamics and enhanced antagonistic sub-ensemble representation of threat-related behaviors, and enhanced threat-related location representation, were also observed. Lastly, cannabinoid administration increased the proportion of SOM neurons exhibiting multidimensional representation of threat-related behaviors and behavior-location conjunction. While cannabinoid receptor activation ex vivo suppressed excitatory inputs to SOM neurons, our data suggest preferential suppression of local GABA release subserves cannabinoid activation of CeA SOM neurons. These data provide insight into how cannabinoid-mediated presynaptic suppression transforms postsynaptic population dynamics and reveal cellular mechanisms by which cannabinoids could affect threat-reactivity.

## INTRODUCTION

Plant-derived cannabinoids including tetrahydrocannabinol and synthetic analogs are widely used for medicinal and recreational purposes and exert broad-ranging behavioral and physiological effects on mood, anxiety, memory, appetite, and sleep via activation of cannabinoid type 1 (CB1) receptors in humans and rodents^1^. CB1Rs are G^i/o^-coupled GPCRs that are widely distributed within the brain and spinal cord and expressed primarily on synaptic nerve terminals and to a lesser degree on postsynaptic membranes, astrocytes, and cellular organelles^2–4^. At synaptic terminals, activation of CB1Rs inhibits neurotransmitter vesicle release and mediates short- and long-term synaptic depression via multiple intracellular signaling pathways, while activation of CB1Rs on astrocytes may lead to synaptic potentiation via release of gliotransmitters^2,3^. Within cortical-like brain regions, CB1 is expressed on a small subset of GABAergic interneurons that form perisomatic contacts with pyramidal neurons and exhibit prominent asynchronous GABA release^5,6^, and more uniformly expressed at low levels within cortical pyramidal neurons within limbic areas, where they are trafficked to axon terminals^5,7^. Subcortically, CB1Rs are expressed in GABAergic neurons of striatal-like regions including the nucleus accumbens and central amygdala (CeA) where they inhibit GABA release locally and in terminal regions^8–11^. Despite these anatomical and synaptic data, how CB1-mediated synaptic suppression ultimately affects neuron population activity, stimulus representation, and dynamics is not well understood.

Of the broad-ranging behavioral effects of cannabinoids, modulation of anxiety and stress-reactivity are prominent and well-described^12–15^. Specifically, while tension and anxiety relief represent major reasons for cannabis use, paradoxical dose- and context-dependent increases in anxiety and panic are also well-documented and some studies have suggested associations between cannabis use and the development and worsening of anxiety disorders^13,16–21^, especially in heavy cannabis users^21^. From a mechanistic perspective, early investigations revealed strong activation of stress-reactive brain regions after cannabinoid administration. Specifically, cannabinoids increase Fos protein expression in the central amygdala (CeA)^22–24^, a key neural substrate for anxiety, emotional learning, and stress-reactivity. Cannabinoid-induced Fos expression in CeA neurons appears synergistic with concurrent stress exposure^23,24^, supporting the notion that CeA may be a relevant substrate for cannabinoid-stress interactions. Accordingly, CB1Rs are heavily expressed in the CeA where they inhibit both glutamate and GABA release onto CeA neurons^8^; however, how these synaptic effects lead to CeA activation and affect CeA population dynamics in response to stress is not known. CeA neurons are heterogenous and express a variety of genetic markers demarcating functionally distinct and antagonistic CeA neuron subpopulations regulating defensive responses^25–27^. Importantly, CeA neurons expressing somatostatin (SOM) are highly stress-responsive and critical for innate avoidance and aversive emotional-learning^28–32^. Taken together, these data suggest activation of CeA SOM neurons could be relevant to the anxiogenic effects of cannabinoids, but how cannabinoids regulate the activity of CeA SOM neurons is not known.

Here, we utilized in vivo single-cell calcium imaging combined with behavioral pharmacology to elucidate the effects of cannabinoid administration on CeA SOM neuron activity and ex vivo optogenetic electrophysiological and slice calcium imaging approaches to elucidate the synaptic mechanisms underlying these effects. Our data show that cannabinoid administration strongly increases the spontaneous activity of CeA SOM neurons and alters neuronal population dynamics associated with acute threat-related behavioral responses. Our data suggest suppression of GABA release from local afferents onto CeA SOM neurons underlies cannabinoid effects on CeA SOM neurons. These data provide insight into how cannabinoid-mediated synaptic suppression transforms postsynaptic population dynamics associated with threat investigation and reveal new cellular mechanisms by which cannabinoids could affect threat reactivity. Elucidating neural mechanisms by which cannabinoids affect stress reactivity and adaptation could ultimately provide important insights into the interactions between cannabis use and anxiety disorders.

## RESULTS

### The cannabinoid agonist CP55940 increases CeA SOM activity in vivo

To begin to elucidate the effects of cannabinoids on CeA SOM neuron activity, we expressed a Cre-dependent GCaMP7f and implanted a 0.66 mm diameter GRIN lens into the CeA of SOM-Cre mice (**Fig. 1 a-b**). Seven mice were injected with vehicle on day 1 and the cannabinoid receptor agonist CP55940 (0.5 mg/kg) on day 2, and calcium transients were compared between pre- and post-treatment conditions in the homecage during a 10-minute recording session. As expected, CP55940 treatment reduced overall locomotor activity (**Fig. S1 a**) and caused a significant increase in the number of active CeA SOM neurons (**Fig. 1 c-e**). Quantification of calcium transients revealed an increase in the AUC per event, but not frequency or amplitude, post-CP55940 treatment (**Fig. 1 f**); effects which normalized 24h after treatment (**Fig. S1 b**). To examine the effects of cannabinoid administration on SOM network properties, we analyzed changes in the degree of correlated activity between CeA SOM neurons and found, in general, sparsely correlated activity (**Fig. S1 c**) with the post-CP55940 distribution significantly shifted toward negative Pearson R values (**Fig. 1 g-h**) and showing lower network centrality (**Fig. 1 i**). We next examined the extent of positive and negative connectivity (defined as correlated activity with a threshold of R > 0.3 or R< -0.3) and found a higher probability of both greater positive and greater negative connection degrees (*k*) post-CP55940 treatment, with substantially greater effect on negative connectivity post-CP55940 (**Fig. 1 j-k**). Despite the shift towards greater values of *k^+^*, positive connection density was reduced and sparsity increased (**Fig. 1 l**), which could be attributed to the fact that the increase in positive connections fall short of keeping up with the number of possible connections due to increase in the number of neurons that are active after CP (**Fig. S1 c**). Conversely, we observe increased connection density and reduced sparsity of negative correlations, on a per mouse basis (**Fig. 1 m**). A similar pattern was observed when analyzing only a subset of neurons, those that were registered during both pre- and post-CP55940 sessions (**Fig. S1 d-e**). Lastly, we analyzed the connectivity of hub neurons, defined as cells with the largest number of connections (>1 SD of mean connections), within and between cellular modules, and found a reduction in the extramodular participation coefficient post-CP55940 treatment (**Fig.1 n-p**). No drug effects were observed on a community structure statistic, quantified as modularity, or the participation coefficient of negative hubs (PC -k) (**Fig. S1 f-g**). Furthermore, while the number of positive connections of positive hubs was not changed, negative connections between hubs and other neurons were increased after CP55940 (**Fig. 1 q**). No changes in any parameter were seen between pre- and post-vehicle treatment (**Fig. S2**). These data indicate cannabinoid administration increases overall activity of CeA SOM neurons, manifested as transients with larger AUCs, and these events tend to show greater degree of negative correlation. In concert, the high degree ‘hub’ neurons show a relative increase in within-module connectivity and increases in number of negative connections with other cells.

**Fig 1.**
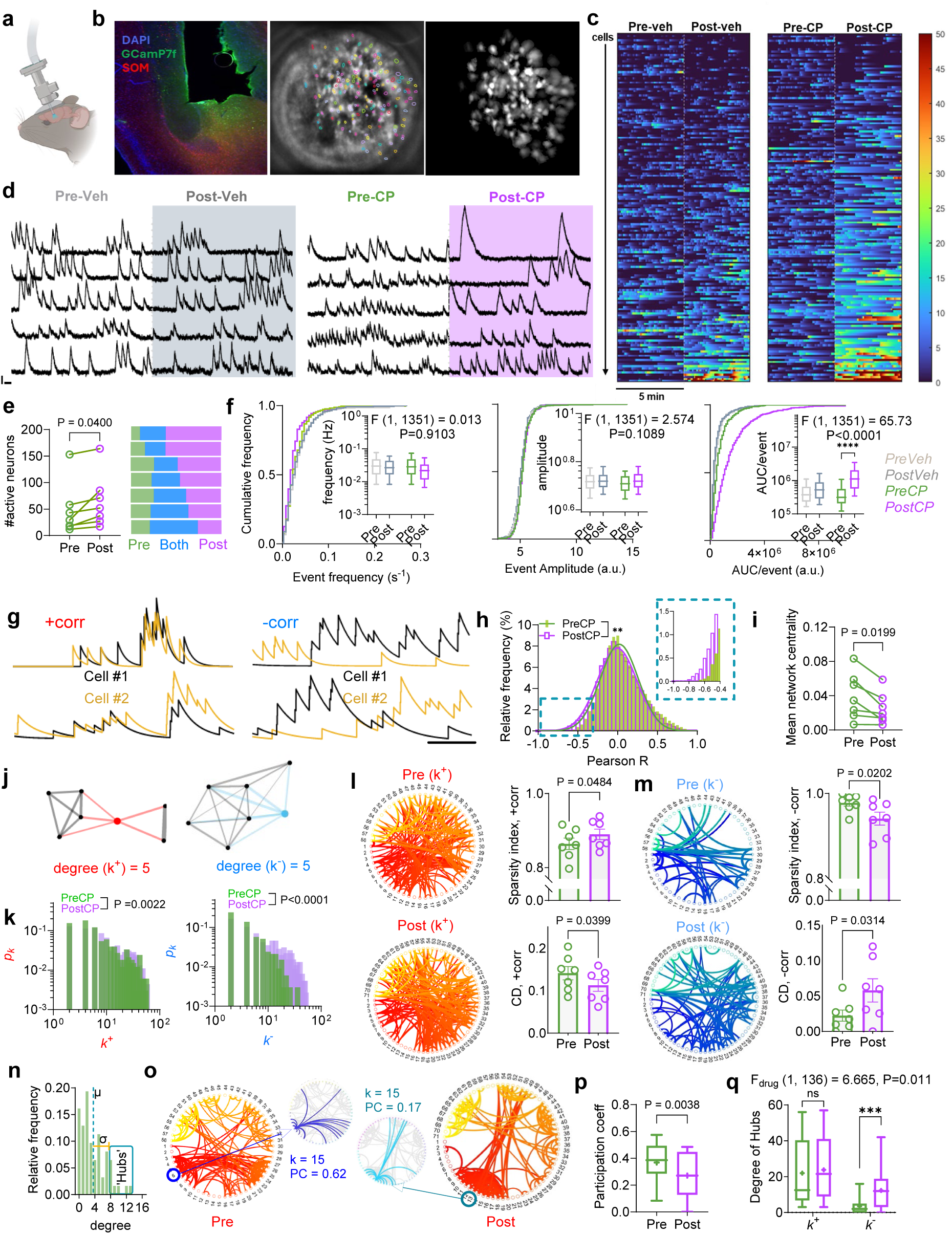
The cannabinoid agonist CP55940 increases CeA SOM activity in vivo. **a)** Schematic diagram of head-mounted miniature microscope for single-cell calcium imaging in freely moving mice. **b)** Representative photomicrograph of CeA GRIN lens placement and GCaMP7f expression in SOM:tdTom-expressing mice and example spatial maps of extracted cells. **c)** Heatmap representing temporal profiles of ΔF/F values over 5 minutes for pre-post-Vehicle sessions (left) and pre-post-CP55940 (CP) sessions (right). **d)** Example traces of spontaneous Ca^++^ transients during the four sessions as in (c) from registered cells from both sessions (scale bar: x = 30s, y=10σ (ΔF/F). **e)** Number of active cells pre-CP and post-CP recorded from N=7(1f,6M) mice (left; P value for paired t-test t=2.612, df=6). Number of cells that were active during the pre, post, and both session per animal (right). **f)** Cumulative distribution and box plots of frequency, amplitude, and AUC per event of extracted Ca^++^ traces (F and P values for 2-way ANOVA followed by Tukey’s multiple comparisons test; ****P<0.0001). Box plots indicate median, interquartile range, and 10th to 90th percentiles of the distribution. **g)** (left) Two representative sets of calcium traces representative of positively (left) and negatively correlated activity (right) (Scale bar: 100 s. **h)** Distribution of Pearson’s correlation coefficients (R) for all simultaneously recorded pairs of neurons from all mice. Inset shows greater relative frequency for negative Pearson’s R values in the post-CP condition. **i)** Mean network centrality is reduced post CP (P value via paired t-test t=3.148, df=6). **j)** Schematic networks of 7 positively (left) and negatively connected nodes (right). Nodes highlighted in red and blue in each network have a degree (k) of 5. (**k)** Degree (k) distributions of positive (defined as Pearson’s R > 0.3) and negative connections (defined as Pearson’s R< -0.3) pre- and post-CP (P values via KS test). **l)** Representative networks of +corr nodes pre- and post-CP administration. Nodes from one mouse are organized at the circumference of the circle and presence of a connections is depicted by curved lines inside the circle. Sparsity index (P value via paired t-test t=2.471, df=6) and connection density (P value via paired t-test t=2.614, df=6) for of networks constructed from +corr connections. **(m)** Same as (l) but for -corr correlations with sparsity index (P value via paired t-test t=3.134, df=6), connection density (P value via paired t-test t=2.793, df=6). Representative network is from the same mouse, but now depicts -corr connections. **n)** Relative distribution of degrees of nodes from all mice with mean (µ) and standard deviation (σ) to categorize a subset of nodes >1 σ from the mean as hubs. **o)** Same representative networks as (l) but depicting community structures via reorganization of nodes around the circumference. Both node 5 (pre-CP) and node 13 (post-CP) are hub neurons and have degrees of 15 (k=15). However, whereas Node 5 has connections within its community and with members of other communities, node 13 has most of its connections within its community. This difference is reflected as a lower Participation Coefficient (PC) post-CP; 0.62 vs. 0.17. **p)** PC values of all hubs from all mice (P value via t-test, t=2.955, df=108). **q)** Comparison of the overall number of positive (+k) and negative (-k) degrees of all hubs. Whereas the number of positive connections between hubs and other nodes is similar pre- and post-CP, the number of negative connections between hubs and other nodes significantly increases post-CP (F and P values via 2-Way ANOVA followed by Sidak’s multiple comparisons test, ***P=0.0003).

### Cannabinoid modulation of SOM neural dynamics associated with threat investigation

Since neural activation of CeA neurons by cannabinoids is synergistic with environmental stress^23,24^, CP55940 increases CeA SOM neuron activity in vivo under baseline conditions, and SOM neurons are activated by stressful experience^28,29^, we hypothesized that CP55940 would modulate threat-induced activation of SOM neurons. To test this hypothesis, we exposed mice to the predator odor analog 2MT^33^, as we have previously described^34^, in a novel cage after vehicle or CP55940 treatment 1 day apart. We analyzed three distinct behavioral motifs during a 20-minute 2MT session; approach, flee, and freezing behavior, as well as time near and far from odor zone (**Fig. 2 a**). Importantly, behavioral responses to repeated 2MT exposure at this re-test interval did not differ in control mice (**Fig. S3 a-b**). Behaviorally, CP55940 shortened the duration of odor investigation, increased freezing, and reduced the number and velocity of approach-flee events (**Fig. 2 b-c** and **Fig. S3 c-d**). We next analyzed neural activity associated with approach, flee, and freezing behavior under vehicle and CP55940 conditions. Consistent with our homecage data, the total number of active cells was higher after CP55940 relative to vehicle treatment (**Fig. 2 d-f**). The proportions of neurons showing increases or decreases in activity (Z-score >±2 SD from baseline) did not differ for approach or flee behavior, but the proportion of activated cells was significantly higher after CP55940 during freezing bouts (**Fig. 2 d-f**). AUC analysis revealed larger magnitude increases and decreases in neural activity for cells exceeding the statistical threshold for pre- and post-approach behavior and for post-flee behavior, without changes during freezing (**Fig. 2 g-i** and quantified in **Fig. S3 e**). Furthermore, we found neural dynamics associated with approach could be used to decode drug states with a high degree of accuracy relative to both shuffled data and neural dynamics associated with either flee or freezing behavior (**Fig. 2 j-l**). Lastly, we determined the effects of CP55940 on the proportion of neurons showing different categorical combinations of response profiles across different behaviors and found significantly larger proportions of neurons that showed antagonistic responses profiles to approach vs. flee and approach vs. freeze, but not flee vs. freeze (**Fig. S4**). These data reveal the presence of SOM sub-ensembles responding antagonistically to specific behaviors (i.e. approach) and across different behaviors (approach and freeze).

**Fig 2.**
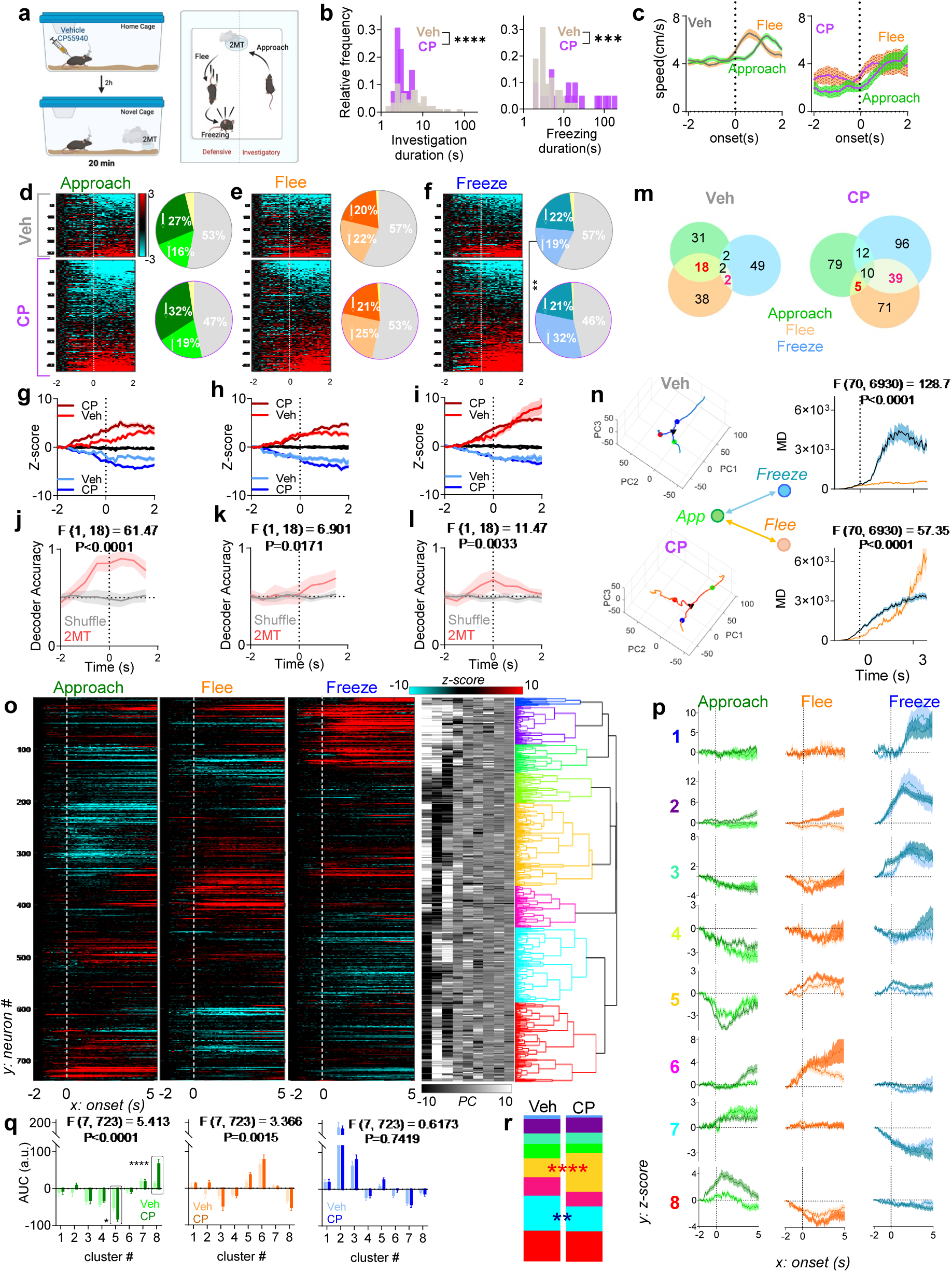
Cannabinoid modulation of SOM neural dynamics during threat investigation. **a)** Schematic diagram of behavioral paradigm. Mice were placed in a cage containing 2MT 2 hours after receiving Veh or CP injection in their home cage and neural activity was quantified during approach, flee, and freezing bouts. **b)** Frequency distributions of duration of 2MT investigation bouts and freezing bouts. CP-treated mice spend significantly less time investigating 2MT per approach bout (KS test, D= 0.6279, ****P=0.0003) and have longer bouts of freezing (KS test, D= 0.5286, ***P=0.0010) Data from N=4 mice (1f,3M). **c)** Speed plots for approach and flee bouts after Veh (n_(Approach)_=129, n_(Flee)_ = 134, left) or CP (n_(Approach)_=14, n_(Flee)_= 17, right). **d)** Heatmaps of Z-scored GCaMP signals of all cells averaged over all approach bouts during Veh (top) or CP (bottom) conditions. Pie charts show the proportion of cells increasing (upward arrow), decreasing (downward arrow) or remaining unresponsive during approach (gray); all defined with a threshold of 2 SD above or below baseline (-2 to -1.5 s). A small fraction of cells (yellow) shows biphasic response. **e)** Same as (d), for flee bouts. **f)** Same as (d), for freezing bouts (+) Responsive cells: χ = 14.24, **P = 1.61e-4, Chi squared test. **g)** Mean Z-score time course for positively modulated (red, Veh: n=52, CP: n=103), negatively modulated (blue, Veh: n=82, CP: n=162), and non-responsive (black) cells time-locked to initiation of approach (time (s) = 0). **h)** Mean Z-scored response of positively modulated (in red, Veh: n=60, CP: n=124) and negatively modulated (in blue, Veh: n=56, CP: n=103) cells time-locked to start of flee. **i)** Mean Z-scored response of positively modulated (in red, Veh: n=55, CP: n=156) and negatively modulated (in blue, Veh: n=62, CP: n=103) cells time-locked to start of freezing. **(j-l)** Accuracy of linear decoder over time in identifying drug condition (vehicle vs. CP) based on neural data associated with approach, flee and freezing bouts relative to shuffled matrix (F and P values via 2-Way ANOVA). **m)** Venn diagrams showing overlap in the number of positively responsive cells (Approach ∩ Flee) - (Approach ∩ Flee ∩ Freeze) Veh=18, CP=5, χ = 24.88, P = 6.10e-7; (Freeze ∩ Flee) - (Approach ∩ Flee ∩ Freeze) Veh=2, CP=39, χ =14.61, P = 1.32e-4. **n)** iPV during approach, flee, and freeze bouts (Veh top left and CP bottom left) projected in PC 1-3 space. Right, time course of Mahalanobis distance (MD) between approach-freeze and approach-flee trajectories under vehicle (top) and CP (bottom) conditions. **o)** Heatmaps of Z scores in response to approach, flee and freeze, sorted according to cluster participation based on hierarchical clustering (of 9 PCs which explain 90% variance) with associated dendrogram. **P-q)** Mean responses of each cluster separated by Veh or CP treatment and their corresponding AUCs (q) plotted for approach, flee and freeze bouts (F and P values for AUC data via 2-Way-ANOVA followed by Tukey’s multiple comparisons test *p=0.047, ****p<0.0001) **r)** Proportions of cells within each cluster after vehicle or CP treatment, respectively; Cluster 1: n=7, n=8; Cluster 2: n=26, n=49; Cluster 3: n=19, n=36; Cluster 4: n=26, n= 31; Cluster 5: n=34, n=126; Cluster 6: n=33, n=47; Cluster 7: n=63, n=81 Cluster 8: n=55, n=98. Cluster 5: χ = 18.316, ****P = 0.00002 and Clust7 χ = 5.197, **P=0.023 via Chi Squared test.

Recent studies have shown distinct subpopulations of CeA SOM neurons respond selectively to a variety of sensory stimuli with little overlap in sensory representation^28^. To evaluate the degree of selectivity of SOM neurons in the representation of threat-related behavioral responses, we next assessed the degree of overlap between behavior-representing activated ensembles and found an increase in in the proportion of cells representing both flee and freeze, and approach and freeze, behavior after CP55940 treatment (**Fig. 2 m** and **Fig. S4 b-c**). No alterations in ensemble overlap were found for behavior-representing inhibited cells (**Fig. S5 a**). These data indicate cannabinoid treatment affects CeA SOM neural dynamics associated primarily with threat investigation (i.e. approach toward 2MT odor) via increased magnitude of changes in activity of antagonistic ensembles representing approach behavior, and proportion of cells with antagonistic responses to different behaviors (i.e. approach vs. flee and approach-vs. freeze). However, these conclusions are based on the statistical categorization of neurons into antagonistic ensembles (i.e. increased or decreased Z-score of >±2 SD). We next wanted to confirm these findings using a complementary approach. Specifically, we first combined all cellular data from vehicle and drug conditions across all three behaviors and performed temporal dimensionality reduction followed by hierarchical clustering (**Fig. 2 o**). Visualization of full temporal data after clustering revealed largely segregated ensembles showing antagonistic activity patterns across all three behaviors (i.e. cluster 5-decreases to approach and increases to flee, and cluster 8-increases to approach and decreases to flee) confirming the presence antagonistic sub-ensembles representing distinct aspects of threat-related behavior exist within the larger CeA SOM neuron population. Next, we separated cells from each cluster based on treatment and compared the AUC between vehicle and drug-treated cells (**Fig. 2 p-q**). This analysis revealed that CP55940-treated cells showed larger AUC for both increases and decreases in activity specifically associated with approach behavior (Clusters 5 and 8) (**Fig. 2 q**). The proportion of cells from CP55940-treated mice was increased within cluster 5 and decreased in cluster 8 (**Fig. 2 r**). These data support the notion that CP55940 increases the magnitude of antagonistic ensemble responses specifically for approach-related behavior and suggests cannabinoids amplify CeA SOM neuron responsivity to threat investigation.

We also assessed the effects of CP55940 on the distance between population vectors associated with each behavior by first performing dimensionality reduction in the pooled activity space and then measuring Mahalanobis distance (MD) for behavior pairs (see methods). When comparing approach vs. flee, MD increased after CP55940, suggesting a more distinct representation of these diverging behaviors after cannabinoid administration (**Fig. S6 a-b**). In contrast, MD was smaller after CP55940 treatment when comparing flee vs. freeze and approach vs. freeze (**Fig. S6 c-f**). We next analyzed the relationship between population vectors when the activity space was confined to each treatment (**Fig. 2 n**). Vehicle treatment was associated with greater divergence of MD between approach and freeze, while with CP55940, MD of both freeze and flee with reference to approach increased steadily (**Fig. 2 n**). Additionally, after vehicle treatment, freeze associated trajectory lengths were significantly longer compared to approach and flee, whereas after CP55940, flee trajectories were longer compared to the other two behaviors (**Fig. S5 b**). Taken together, this shows that neuronal population vector trajectory undergoes significant alterations both in the distance it traverses and in temporal dynamics of the relationship between vectors representative of each behavioral state.

### Cannabinoid modulation of location-behavior co-representation in CeA SOM neurons

In addition to behavioral responses to 2MT, mice also changed their preferred location within the cage to maximize distance from the odor area, and CP55940 treatment increased this avoidance behavior (see Fig. S3 c). Accordingly, we next determined whether CeA SOM neurons were able to represent threat-related location and if this representation was altered by cannabinoid administration. Furthermore, since our data above indicate cannabinoid treatment increased multidimensional representation of threat-related behavior (see Fig. 2 m), we also examined the degree of co-selectivity between location representation and mobility as an index of conjunctive location-behavior representation (i.e. multidimensional location-behavior representation).

As expected from our earlier analyses, CeA SOM neurons showed multiple distinct phenotypes including cells active during periods of immobility, in the near odor zone, and cells that show both mobility and near odor zone activity, for example (**Fig. 3 a-b**). To systematically analyze cellular activity associated with location and mobility, we Z-scored cellular activity to the entire 20-minute session and defined periods of neuronal activation as periods when the Z-score exceeded 1.96. We next analyzed the probability of cellular activation during two opposing epochs of location (near odor vs. far odor) and behavior (mobility vs immobility). Examination of cell activation probability distributions revealed CP55940 increased cellular activation during both near and far from odor locations (**Fig. 3 c**). Similarly, CP55940 increased the probability of cell activation during both mobile and immobile periods (**Fig. 3 d**). These data suggest cannabinoid administration increases population activation during epochs of interest consistent with earlier results showing general increases in SOM activity after CP55940 treatment, but do not reveal whether individual cells show changes in selectivity magnitude for location or behavior after drug treatment. To explicitly test this, we generated a per-cell selectivity index (see methods) for near vs. far zone and mobility vs. immobility. Indeed, both vehicle and CP55940-treatment exhibited cell subsets with selectivity for near vs. far zone, and CP55940-treament showing differences for mobility vs. immobility, relative to scrambled control data sets (**Fig. 3 e and g**). To quantify differences in cellular selectivity between drug and vehicle conditions, we compared the selectivity index of statistically selective cells (those with a selectivity index > ± 2SD from their corresponding shuffled distribution). This analysis revealed cannabinoid administration increased the magnitude of selectivity for statistically far-selective neurons and both mobility- and immobility-selective cells. For near selective neurons, a small group of highly selective neurons were present, which were not seen after vehicle treatment (**Fig. 3 f and h**). We next examined whether cannabinoid administration affected the degree of conjunctive location-behavior representation of CeA SOM neurons. We first plotted the selectivity index for near vs. far and mobile vs. immobile periods for each neuron together. These data clearly show cells from CP55940-treated mice exhibited high degrees of co-selectivity for both near and far selectivity with mobility, relative to vehicle treatment (**Fig. 3 i**). We additionally examined the proportion of cells showing conjunctive selectivity between mobility and immobility with near-odor representation, and far-zone representation (**Fig. 3 j**). This analysis revealed significant increases in multidimensional location-behavior representation, and a reduction in unidimensional location representation i.e. selective for only near or only far locations, after CP55940 treatment. These data suggest cannabinoids increase the magnitude of selectivity of cells representing threat-related location and behavior and increase the degree of conjunctive location-behavior representation of CeA SOM neurons.

**Fig 3.**
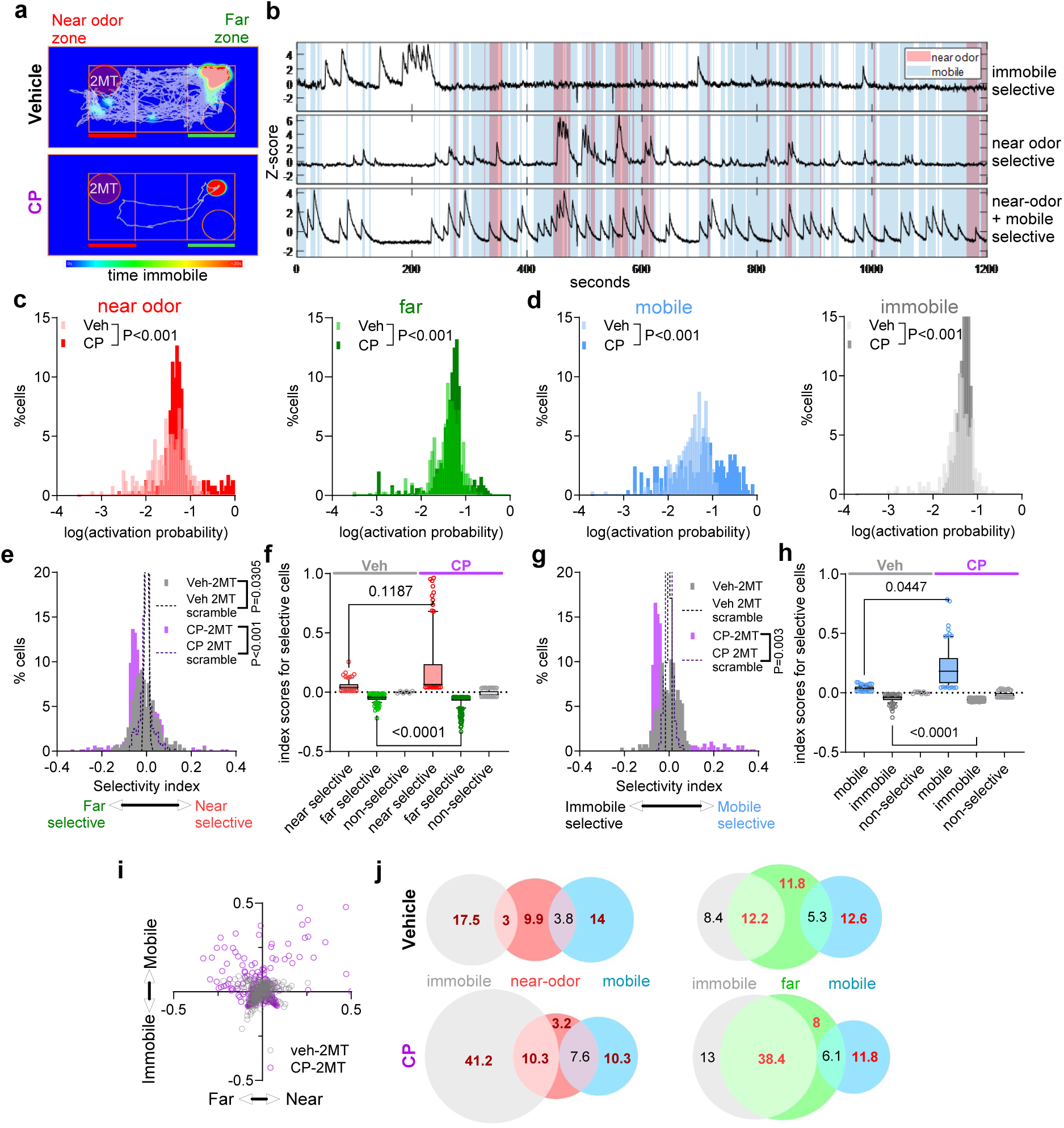
Cannabinoid treatment enhances threat-related location and behavior representation. **a)** Representative locomotion trajectory (grey lines) and immobility (heatmap overlay) of a mouse during 2MT exposure post-vehicle (top) and post-CP (bottom) administration. **b)** Representative Z-scored GCaMP signals from cells selective for immobility (top), near-odor zone occupancy (middle) and, both near-odor and mobile bouts (bottom). Cells were considered selective if their respective index values exceeded ± 2 SD of scrambled dataset for either location or behavior (see e and g). **c)** Relative distribution of cell activation probabilities (see methods) while mice occupied the near odor zone (left) or far zone (right). **d)** Same as (c), for mobile and immobile bouts. **(e)** Cell activity selectivity index histogram for zone occupancy. Positive values indicate preferential activation during near-odor zone occupancy while negative values indicate preferential activity during far-zone occupancy. Scrambled selectivity distributions for vehicle and CP conditions shown in broken lines. P values via KS test. **f)** Selectivity index values for zone-occupancy selective (index values > ± 2 SD of corresponding scrambled dataset) and non-selective cells after vehicle or CP administration (P values via Kruskal Wallis test followed by Dunn’s multiple comparisons test). **g)** Cell activity selectivity index histogram for mobility. Positive values indicate preferential activation for mobile bouts while negative values indicate preferential activity during immobile bouts. **(h)** Selectivity index values of mobile and immobile selective cells (P values via Kruskal Wallis test followed by Dunn’s multiple comparisons test). **i)** Summary scatter plot of mobility and zone occupancy selectivity indices for all cells depicting CP-induced emergence of cellular subsets with elevated multidimensional selectivity; high selectivity values for both dimensions (mobility and location). **j)** Venn diagrams depicting percentage of immobility/ mobility selective cells and their overlap with location selective cells: near odor zone (left)/ far zone (right). There is an increase in proportion of cells with dual behavior-location selectivity and reduction in only location selectivity after CP administration. Significant groups differences are denoted in red: Immobile – (Immobile ∩ Near): Veh=17.5%, CP=41.2%, χ = 15.75, P=7.2e-05; (Immobile ∩ Near): Veh=3%, CP=10.3%, χ = 5.461, P=0.019; Near - (Immobile ∩ Near) - (Mobile ∩ Near): Veh=9.9%, CP=3.2%, χ = 30.43, P=3.5e-08; Mobile – (Mobile ∩ Near): Veh=14%, CP=10.3%, χ = 13.89, P=0.00019; (Immobile ∩ Far): Veh=12.2%, CP=38.4%, χ = 25.75, P=3.9e-07; Far - (Immobile ∩ Far): Veh=11.8%, CP=8%, χ = 14.14, P=0.00017; Mobile - (Mobile ∩ Far): Veh=12.6%, CP=11.8%, χ = 6.35, P=0.0117, Chi-squared test).

### Cannabinoid modulation excitation-inhibition balance onto CeA SOM neurons

Our data thus far indicate cannabinoid administration increased overall SOM neuron activity, neuronal dynamics associated with threat-related behavioral reactivity, and location-behavior co-representation. To gain insight into the synaptic mechanisms underlying cannabinoid activation of SOM neurons, we utilized whole-cell patch-clamp electrophysiological and slice calcium imaging approaches. As expected and consistent with our previous published work^8^, CP55940 reduced excitatory and inhibitory synaptic currents onto CeA SOM neurons, while CB1 receptor blockade did not affect either revealing a lack of endogenous cannabinoid tone at these synapses (**Fig. 4 a-e**). Given these data provided little insight into how CP55940 could ultimately increase SOM neuron activity in vivo, we next examined the effects of CP55940 on distinct excitatory afferents to CeA SOM neurons. Specifically, we analyzed dorsal medial thalamic (dMT) and parabrachial nucleus (PBN) afferents given the critical role of PBN-dMT pathways in the regulation of predator avoidance^35,36^ and their strong synaptic connectivity with CeA neurons^37^ (**Fig. 4 f-g**). Optogenetic stimulation of dMT and PBN afferents elicited monosynaptic excitatory and di-synaptic inhibitory currents in CeA SOM neurons (**Fig. 4 h-i**). Subsequent current-clamp recordings revealed distinct properties of postsynaptic potential responses to dMT and PBN stimulation (**Fig. S7 a-c**), and differential temporal integration (facilitation for dMT and depression for PBN) and recruitment of disynaptic GABAergic transmission (**Fig. S7 d-e**). Both inputs showed greater inhibitory current amplitude and area and increased I/E ratios (**Fig. 4 h-i** and **Fig. S8 a-c**), and the I/E ratios were substantially larger for a subset of PBN stimulated cells relative to the dMT. Interestingly, PBN excitatory and disynaptic inhibitory responses showed greater presynaptic release probability than dMT responses (**Fig. S8 d**), further suggesting PBN inputs may recruit distinct disynaptic GABAergic neurons for feedforward inhibition. Application of CP55940 reduced excitatory and disynaptic inhibitory currents elicited by dMT stimulation (**Fig. 4 j-k**), but preferentially inhibited disynaptic GABA release, over glutamate, upon PBN stimulation (**Fig. 4 l-m**). Indeed, the I:E ratio was also reduced post-CP55940 for PBN, but not dMT stimulation (**Fig. 4 n-o**), and direct comparison of CP55940 effects on both afferent types revealed significant interaction with reduced suppression of excitation from PBN afferents relative to dMT afferents by CP55940 (**Fig. 4 p**). Effects of CP55940 on glutamate and GABA were presynaptic, as an increase in the paired-pulse ratio (EPSC_2_/EPSC_1_) was observed for both (**Fig. 4 q** and **Fig. S8 e**). These data suggest cannabinoids could activate CeA SOM neurons via a disinhibition mechanism exerted preferentially upon the brainstem, over thalamic, afferent activation onto SOM neurons.

**Fig 4.**
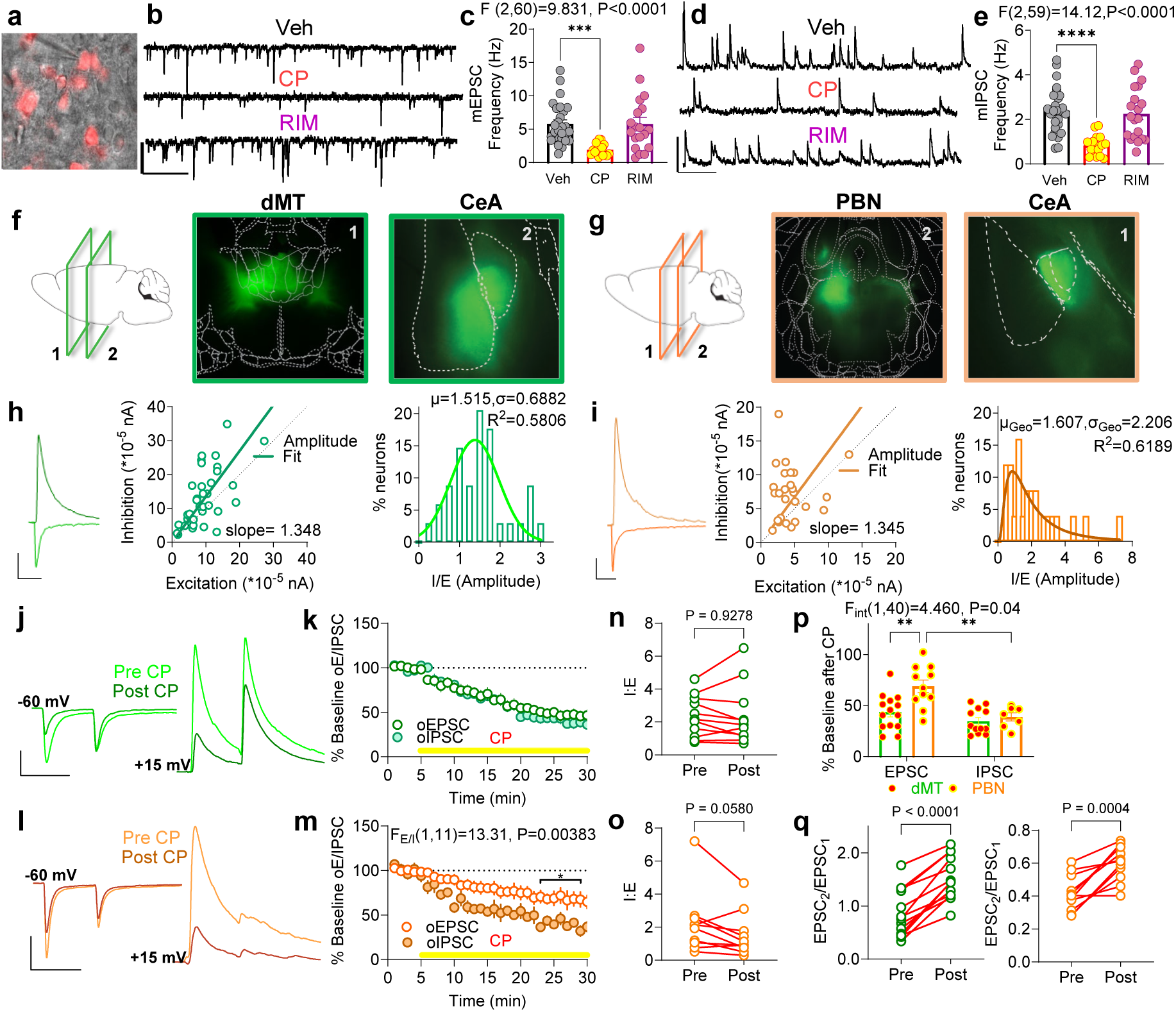
Cannabinoid modulation of excitation-inhibition balance onto CeA SOM neurons. **a)** Representative DIC 40X brightfield image of CeA slice with recording pipette and td-tomato expressing CeA SOM neuron. **b)** Representative mEPSC recording traces (Scale bar: 40 pA, 500 ms) **c)** Effects of CP55940 or Rimonabant on mIPSC frequency (Ordinary one-way ANOVA, Dunnet’s multiple comparison, ***P=0.0003). **d)** Example traces of mIPSC recording (Scale bar: 50 pA, 500 ms) **e)** Effects of CP55940 or Rimonabant on mIPSC frequency (Ordinary one-way ANOVA, Holm-Sidak’s multiple comparison, ****P<0.0001). **f)** Schematic of approximate location of coronal planes 1 and 2 imaged at 4X, with ChR2 expression in the dMT injection site (#1) and expression of ChR2 terminals in slice containing the CeA (#2).**g)** Schematic of approximate location of coronal planes 1 and 2 imaged at 4X, with ChR2 expression in the PBN injection site (#2) and expression of ChR2 terminals in slice containing the CeA (#1) **h)** Representative EPSC and IPSC recorded from CeA SOM neurons in response to optical stimulation of dMT inputs (scale bar: 200 pA, 50 ms). XY scatter plot of peak EPSC and IPSC amplitudes in the same cells, with linear regression. Summary of I/E ratio, which was best fit by a gaussian model. **i)** Same as g for PBN input stimulation (scale bar: 200 pA, 50 ms). For this input, the frequency distribution of IE ratios was best fit by lognormal model. **j)** Example current traces of optically evoked pairs of EPSCs (at -60mV) and IPSCs (at +15 mV) recorded before and after CP wash-on (scale 500 pA, 50 ms,). **k)** Time course of bath application of CP on EPSC and IPSC amplitudes upon optical stimulation of dMT-CeA synapses. **l)** Example traces of optically evoked pairs of EPSCs and IPSCs recorded before and after CP wash-on (scale 200 pA, 50 ms). **m)** Time course of bath application of CP on EPSC and IPSC amplitudes upon optical stimulation of PBN-CeA synapses showing a significant greater reduction in IPSC amplitude (F and P values from 2-Way ANOVA followed by Holm-Sidak’s post hoc, *P<0.05). **n-o)** Effect of CP on I/E ratio upon stimulation of dMT-CeA (top) PBN-CeA (bottom) synapses (P values via pared t-test t=0.0929 df=10 for dMT, t=2.172 df=9 for PBN). **p**) Summary analysis of the effects of CP on EPSC and IPSC depression at dMT-CeA or PBN-CeA synapses showing asymmetric preferential suppression of disynaptic GABA release at PBN-CeA synapses over dMT-CeA synapses (F and P values via 2-Away ANOVA followed by Tukey’s post hoc test, **P<0.01). **q**) Effect of CP wash-on on EPSC PPR upon optical stimulation of dMT (left) or PBN afferents (right) (P values via paired t-test, dMT: t=8.879, df=12, PBN: t=5.209, df=10).

### Cannabinoid suppression of GABA transmission activates CeA SOM neurons

To explicitly test whether cannabinoid suppression of GABA can disinhibit SOM neuron firing, we first determined the effects of CP55940 on distinct GABAergic synapses onto SOM neurons and found both GABAergic inputs from other SOM neurons and non-SOM-expressing neurons (SOM^-^) synapse onto SOM cells and are inhibited equally by CP55940 (**Fig. 5 a-c**). We then expressed ChR2 in SOM^-^ CeA cells using a Cre-out system and evaluated the ability of SOM^-^ neurons to suppress AP firing of SOM cells in the absence or presence of CP55940 (**Fig. 5 d**). In the absence of CP55940, activation of SOM^-^ neurons robustly suppressed depolarization-induced firing of SOM neurons, an effect reliably attenuated in the presence of CP55940 (**Fig. 5 e**). Similar effects were observed when stimulating SOM^-^ neurons during the peak (-20 to 0 ms) of postsynaptic sinusoidal current injection (**Fig. 5 f-g**). These data indicate that SOM^-^ cells can robustly suppress the firing of SOM neurons and that CP55940 attenuates this suppression through inhibition of GABA release from SOM^-^ to SOM neurons. These data support the notion that cannabinoids can increase SOM neuron activity via a disinhibitory mechanism.

**Fig 5.**
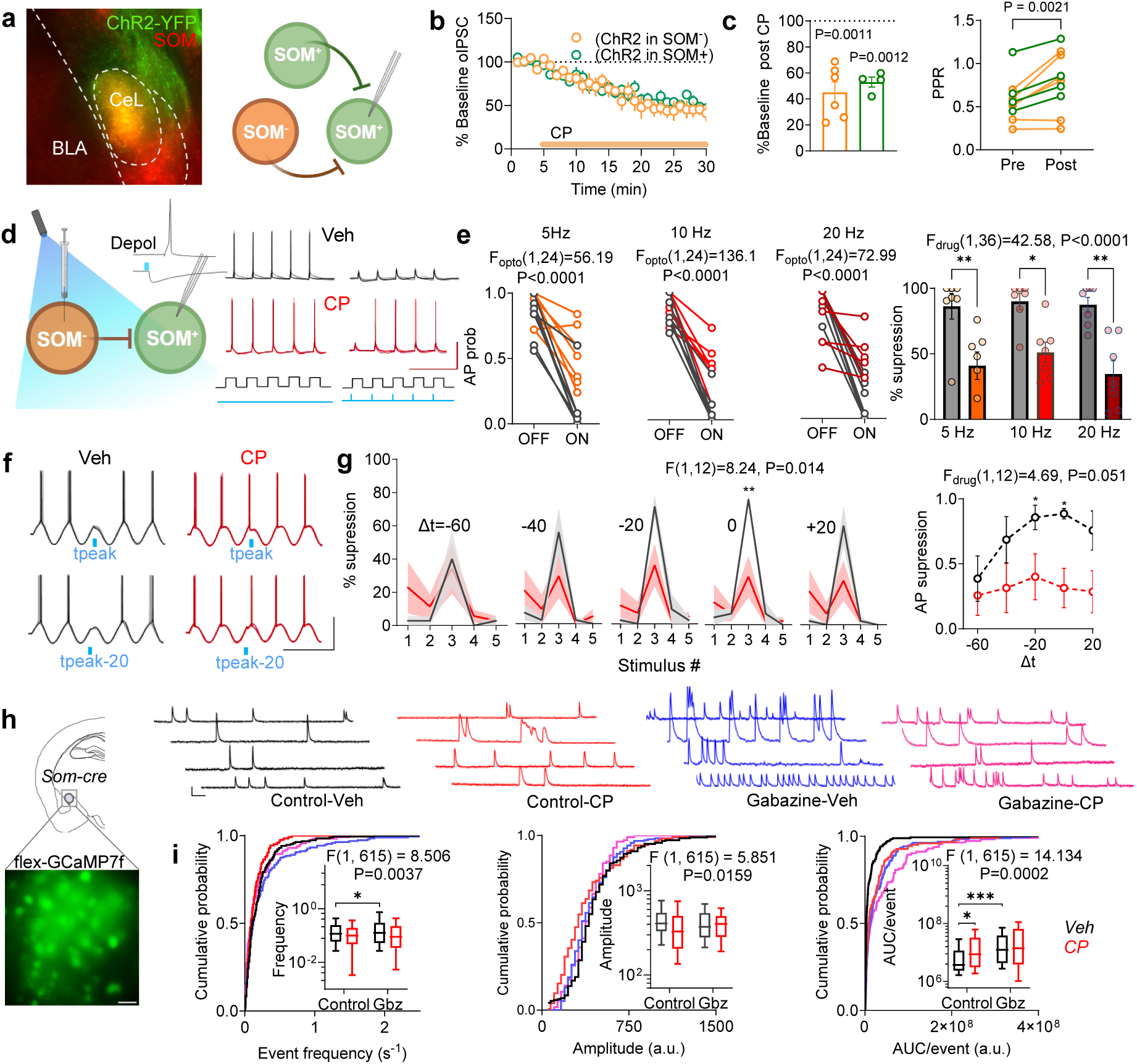
Cannabinoid suppression of local GABA transmission activates CeA SOM neurons. **a)** Representative image of ChR2 expression in tdTom-expressing CeA SOM cells. IPSC recordings were performed from CeA SOM neurons while stimulating terminals from ChR2 expressing terminals arising from either SOM+ or SOM-neurons. **b)** Time course of CP-induced suppression of IPSCs onto CeA SOM neurons from SOM+ or SOM-cells. **c)** Summary of average reductions in IPSC amplitude and PPR in the last 5 mins compared to the 5 min baseline (p values via One sample t-test; SOM^−^: t=6.759, df=5, SOM^+^: t=12.23, df=3). **d)** Schematic description of experimental strategy: APs were elicited by depolarizing current injection in SOM cells. Activation of virally expressed ChR2 in SOM-terminals evokes IPSPs in SOM cells. Example voltage traces showing firing with current pulse at 5 Hz and spike suppression with concurrent presentation of blue light under vehicle (top) and CP conditions (below) Scale bar: 50 mV, 500 ms. **e)** Plots of action potential probability upon 5, 10 and 20 Hz depolarization during light-ON and light-OFF conditions under vehicle and CP conditions. SOM- cells strongly suppressed AP firing in SOM cells. CP application significantly reduced suppression of AP spiking across all frequencies (F and P values via Two-way ANOVA followed by Tukey’s multiple comparison test, *p<0.05, **p<0.01). **f)** Illustrative traces of APs elicited by injection of sinusoidal current waveforms, with concurrent optical activation of SOM^-^ synapses (wave 3) at different times with respect to the peak of the cycle (-60 ms to +20 ms, Scale bar: 50 mV, 500 ms). **g)** Maximal suppression occurred when ChR2 stimulation was delivered near the peak of the sine wave (Δt=0) and the efficacy of suppression dropped steeply as the IPSP moves further from the time of the peak. CP application attenuated light-evoked suppression most effectively at -20 and 0 ms timepoints (F and P values via 2-Way ANOVA followed by Tukey’s post hoc comparisons *p<0.05, **p<0.01). **h)** Representative image showing GCaMP7f expression in SOM cells from CeA slices and example traces of ex vivo Ca^++^ transients under control, CP, Gabazine and Gabazine + CP conditions Scale bar: 2e03 a.u., 2 s. **i)** Cumulative probability and box plots of event frequency, amplitude and AUC/event across the four groups (F and P values via 2-Way ANOVA followed by Tukey’s post hoc comparisons; *p<0.05, *** p<0.001).

To confirm this hypothesis, we examined the changes activity of CeA SOM neurons in acute brain slices using calcium imaging approaches in response to CP55940 in the absence or presence of the GABA_A_ receptor blocker gabazine. Under control conditions, CP55950 increased the AUC/event of CeA SOM like the effects observed in vivo, confirming cannabinoids activate SOM neurons via actions locally in the CeA and do not require intact brain-wide neural circuit dynamics (**Fig. 5 h-i**). Importantly, gabazine alone also increased the AUC/event in a manner similar to CP55940 and occluded any further increase in AUC/event by CP55940. These data provide corroborating evidence that cannabinoids increase CeA SOM activity via a disinhibitory mechanism arising from the suppression of local GABA release in the CeA.

## DISCUSSION

Cannabinoids are well known to trigger anxiety reactions in humans and increase avoidance and threat reactivity in rodents^12,14,15,18^, however, the mechanisms by which cannabinoids affect innate avoidance are not well understood. Importantly, activation of CeA neurons has been suggested to contribute to the anxiety-modulating effects of cannabinoids^23,24^ and CB1 receptor activation modulates glutamatergic and GABAergic synaptic transmission onto CeA neurons^8^. Here we show that the potent cannabinoid agonist, CP55940, robustly increases the activity and alters network dynamics to favor the emergence of antagonistic representational sub-ensembles of CeA SOM neurons, which are highly threat reactive critical for the generation of defensive responses^27–29^. CP55940 increased predator odor-induced threat avoidance, which was associated with significant increases in antagonistic ensemble representation of threat-investigatory behavior. Moreover, the selectivity of threat-related location and multidimensional location-behavior representations were enhanced after cannabinoid treatment. Lastly, our synaptic and ex vivo imaging studies indicate cannabinoid-mediated suppression of local GABAergic transmission contributes to enhanced SOM neuron activity. These data provide insight into how cannabinoid-mediated presynaptic suppression transforms postsynaptic neural dynamics supporting enhanced threat avoidance, suggest cannabinoid activation of SOM neurons as a mechanism underlying anxiogenic effects of cannabinoids, and provide new insights into CeA function by revealing the functional presence of within cell-class antagonistic ensembles representing distinct threat-related behavioral responses.

To gain insight into how cannabinoids affect CeA SOM neuron activity in vivo, we conducted calcium imaging studies followed by a series of network theory-based analyses^38^. Our data reveal four major effects of CP55940 on basal activity of the CeA SOM neuron network: 1) an increase in activity of SOM neurons measured as an increase in the number of active neurons and AUC per calcium event, 2) a shift toward increased negatively correlated population activity, evidenced by a reduction in the density of positive connections and substantial increase in negative connection density, 3) an increase in the ratio of within-module to between module positive connections of hub neurons, and 4) increased number of negative connections between hubs and other neurons in the network. These data indicate that cannabinoids increase the overall activity of the SOM neuron population in vivo and could alter network dynamics to support synchronization of sub-ensembles (via increased within-module correlated activity or network hubs) and promote the generation of antagonistic sub-ensembles (via increase in degree of negative connections of network hubs).

To determine how these basal changes in network dynamics affect SOM neuron representation of threat-related behavior, we examined SOM activity during approach, flee, and freezing responses to predator odor exposure. As expected, CP55940-treated mice exhibited increased avoidance of predator odor, increased freezing and overall immobility^10,39^, and lower approach and flee bouts and reduced speed^39^, relative to vehicle treatment. In line with our analysis of basal network changes induced by CP55940, data using both threshold-based and unbiased clustering approaches revealed the magnitude of changes in activity for antagonistic ensembles was augmented by cannabinoid treatment, an effect which was most robust for threat investigation (odor approach). Indeed, decoder accuracy for drug-state (CP55940 vs. vehicle) was highest for approach-related neural dynamics over flee- or freezing-related dynamics. It is noteworthy that our data reveal neural activity changes prior to movement onset which is consistent with data from upstream BLA neurons^40^, whereas other studies have found CeA SOM neuron activity lags behavior^28^; differences possible due to methodological approaches to capturing behavior onset (i.e. licking vs approach initiation). Similarly, a previous study found little evidence for sensory stimuli-driven SOM neuron inhibition or the presence of antagonistic sub-ensembles in response to distinct sensory stimuli^28^. Indeed, current conceptualizations of CeA function posit the interplay of ensembles of functionally antagonistic and genetically non-overlapping cell types (i.e. SOM vs. PKCδ) ultimately guide behavioral selection via a winner-take-all type of mechanism^25–27,37,41,42^. Evidence for this model exists when considering defensive^25–27,41,42^ and pain-related behavioral responses^43^. Our data challenge this view by presenting compelling evidence of antagonistic sub-ensembles *within* the CeA SOM neuron population itself, with different SOM cells that show either increases or decreases in activity to all three threat-related behaviors analyzed in approximately equal proportions. Antagonistic activity within sub-ensembles was also evident across different behaviors (i.e approach-flee and approach-freeze). Our synaptic data provide further support for this model. Specifically, while demonstration of GABAergic synaptic connectivity between SOM^−^ and SOM+ cells is consistent with previous data and likely supports generation of *between* cell-class antagonistic ensembles, we also detected GABAergic synapses between SOM+ and other SOM+ cells, which could provide a functional basis for generation of *within* cell-class antagonistic ensembles.

In addition to threat-related behavior, we found SOM neurons also represented threat-related location as determined by proximity to odor (near vs. far). We found overall increases in the activation probability distributions and selectivity indices for both near and far active cells after CP55940 treatment relative to vehicle or corresponding scrambled matrix, respectively. Similar effects were observed when analyzing mobility behavior, suggesting that both spatial and behavioral selectivity of SOM neuron activity is enhanced after cannabinoid treatment. These data suggest the possibility that increased behavioral avoidance may be related to augmented neural representation of potentially dangerous and safe locations after cannabinoid treatment.

While simple sensory or behavioral selectivity is a conserved feature of primary cortical areas, multidimensional coding or mixed selectivity appears to be a feature of associative and limbic cortical areas including the prefrontal cortex and basolateral amygdala^44^. For example, prelimbic prefrontal cortical neurons are more likely to be active during specific spatial location-social interaction conjunction than either behavior or location alone^45^, and BLA neurons show multidimensional responses to conditioned stimuli and conditional behavioral responses^46^ and social stimuli^47^. That BLA neurons project to CeA SOM neurons^30,48^ suggests that these representational properties would be transferred to projection targets. Indeed, several pieces of data presented here support this notion. First, the overlap between SOM neurons activated during freezing and flee behavior increases after cannabinoid administration. Second, the proportion of neurons representing both threat-related location (near vs. far) and mobility (mobile vs. immobility) increases after cannabinoid treatment. Third, cannabinoid treatment increases multi-dimensional selectivity, specifically for mobility and location conjunction, over immobility and location conjunction. Several theoretical models could explain these results. For example, since the total number of active CeA SOM neurons is increased by cannabinoid administration, it is possible that newly recruited neurons exhibit an intrinsically higher degree of multidimensionality, and any condition that recruits progressively more SOM neurons will result in a commensurate increase in the degree of multidimensionality. Alternatively, cannabinoid treatment could fundamentally change the representational properties of SOM neurons such that their activity is regulated by a broader range of stimuli. Indeed, cannabinoid-mediated suppression of GABA release onto SOM neurons could serve as a synaptic substrate for both possibilities (see below). Interestingly, these data are consistent with previous experiments demonstrating increases in the overlap in the representation of unconditioned and conditioned sensory stimuli representation in CeA SOM neurons that develop across a Pavlovian learning task^28^. Furthermore, the number of unconditioned and conditioned stimuli-responsive neurons increases across training^28^; both effects are strikingly similar to those observed after cannabinoid administration. Whether these changes are mediated via endogenous cannabinoid release and CB1 receptor activation remains an intriguing possibility to be tested in future work.

We have also aimed to uncover underlying synaptic mechanisms subserving the overall increase in CeA SOM neuron activity. Taken together our data suggest suppression of GABA release onto SOM neurons (from SOM^-^ and other SOM neurons) represents a major mechanism underlying cannabinoid-induced increases in SOM neuron activity. We first examined how PBN and dMT afferents, which have been causally implicated in predator avoidance^35,36^, form excitatory synapses onto CeA SOM neurons and trigger disynaptic GABAergic responses that exhibit different synaptic properties. Importantly, we found that CP55940 suppressed GABA and glutamate release equally from dMT afferents, but CP55940 depressed GABA to a greater extent than glutamate release upon PBN afferent stimulation, suggesting cannabinoids may preferentially disinhibit SOM neurons via an asymmetric synaptic suppression and changes in excitation-inhibition balance. Our observation that PBN I/E ratio follows a lognormal distribution posits that a small but significant proportion of PBN inputs could drive very strong disynaptic inhibition and that CP55940-inducd depression of these GABAergic inputs could have a nonlinear multiplicative effect to increase network excitability^49^. Importantly, suppression of SOM neuron firing by activation of local SOM^-^ CeA neurons is attenuated after CP55940 administration indicating that cannabinoids can disinhibit SOM neuron firing ex vivo. Lastly, we show that blocking GABA_A_ receptors mimics and occludes the increases in AUC per calcium event observed after CP55940 in acute brain slices. These data suggest a primary role of disinhibition in the cannabinoid activation of CeA SOM neurons in vivo.

Several technical considerations and study limitations are also worthy of mention. First, we used a relatively high dose of CP55940 (0.5 mg/kg) to trigger prominent avoidance and maximize detection of changes in SOM neuron activity. We have previously shown doses between 0.3 mg/kg and higher increase Fos expression in CeA neurons^24^ and other cannabimimetic effects including hypothermia and analgesia increase linearly between 0.1 and 1 mg/kg^50^. In an extension of this issue, the profound avoidance observed at this dose resulted in a significantly lower number of avoidance bouts and duration spent in the near odor zone. This difference resulted in an asymmetric number of data points for location and behavioral selectivity measures, for example. However, to avoid biases due to the aforementioned asymmetry, we used the same statistical cutoff for selectivity relative to each group’s scrambled matrix to maximize the validity of our conclusions. Similarly, as expected the speed of approach-flee bouts was significantly lower in CP55940-treated mice^39^, however, the profound neural dynamic changes observed were relatively selective for approach behavior, despite both approach and flee having equally depressed speed relative to vehicle treatment. In this context, future studies examining lower doses and overtly anxiolytic doses (0.001-0.01 mg/kg)^51,52^ on SOM neuron activity would be important. Lastly, how cannabinoids affect other CeA cell types and the effects of cannabinoids on SOM neuron representation of appetitive approach behavior remain to be tested as SOM cells also respond to positively valanced stimuli such as sucrose^28^.

Cannabinoids are well-known to precipitate anxiety and panic reactions, especially at high doses under aversive or novel environmental contexts^12,15,18,19^. Moreover, associations between cannabis use and anxiety disorders are well-documented^13,16–21^. Importantly, a recent large-scale longitudinal study of 12 million subjects suggests cannabis use may precede the development of anxiety disorders in some individuals^21^. While causal links between cannabis use and the development of anxiety disorders are not conclusive, cannabis use associated with an emergency room visit, presumably due to serious adverse effects, was associated with a 3-fold greater risk of subsequent medical visits for an anxiety disorder^21^. Our data presented herein begin to shed light on neural mechanisms subserving potential “dark-side” adverse effects of cannabinoids which are likely to become more prevalent as cannabis use increases nationally. Specifically, we suggest cannabinoids could trigger adverse anxiety and panic reactions via activation of CeA-related circuits and that cannabis-induced long-term adaptations in these circuits could contribute to observed associations between cannabis use and anxiety disorders.

## MATERIALS AND METHODS

### Animals

All experiments were approved by the Northwestern University Animal Care and Use Committee or Vanderbilt University Institutional Animal Care and Use Committees and were conducted in accordance with the National Institutes of Health Guide for the Care and Use of Laboratory Animals. Adult C57BL/6J mice of both sexes were used, and data were pooled from both sexes as no overt sex differences were observed. Mice were group housed (2-4 animals per cage) in a temperature- and humidity-controlled housing facility under a 12 h light-dark cycle with ad libitum access to food and water. Mice were implanted with Gradient Index (GRIN) lenses which were single housed. SOM:Ai14 mice and SOM-Cre heterozygous lines used in this study were obtained by crossing homozygous transgenic line SOM-IRES-Cre (JAX stock #013044) with Ai14 (JAX stock #007914) and wild-type C57BL/6J respectively. All experiments were performed during the light cycle.

### Viruses

For ex vivo circuit mapping of external inputs to the CeA, the following viral injections were performed in SOMAi14 mice: AAV5-CaMKIIα-hChR2(H134R)-eYFP (0.3–0.35 μl, Addgene# 26969-AAV5, titer ≥ 1×10¹³ vg/mL) was injected unilaterally to either dMT or PBN (coordinates noted below). AAV5-EF1a-double floxed-hChR2(H134R)-EYFP-WPRE-HGHpA (Addgene# 20298, titer ≥ 1×10¹³ vg/mL) or AAV2-EF1a-FLEX/OFF-hChR2(H134R)-eYFP (0.25 μl, titer ≥ 1×10¹³ vg/mL, gift from Dr. Penzo lab, NIH, NIMH)^53^ was bilaterally injected in the CeA for mapping local inhibition within CeA. AAV9-syn-FLEX-jGCaMP7f-WPRE (0.35 μl, Addgene# 104492, titer ≥ 1×10¹³ vg/mL) or retroAAV-syn-jGCaMP7f-WPRE (0.35 μl, Addgene# 104488, titer ≥ 7×10¹² vg/mL) was injected unilaterally in SOMAi14 mice for microendoscope recordings. AAV9-syn-FLEX-jGCaMP7f-WPRE (0.4 μl, Addgene# 104492, titer ≥ 1×10¹³ vg/mL) was injected bilaterally in SOM-Cre mice for slice calcium recordings.

### Surgery

At 6-8 weeks of age, mice were anesthetized with 5% isoflurane. The skin on the incision site was prepped with alcohol and iodine after the hair was shaved. The animal was transferred to a stereotaxic frame (Kopf Instruments, Tujunga, CA) and kept under 1-2% isoflurane anesthesia. The skull surface was exposed via a midline sagittal incision and treated with the local anesthetic benzocaine (Medline Industries, Brentwood, TN). For each surgery, a 10uL microinjection syringe (Hamilton Co., Reno, NV) with a Micro4 pump controller (World Precision Instruments, Sarasota, FL) was guided by a motorized digital software (NeuroStar; Tübingen, Germany) to each injection coordinate. The following coordinates were used relative to Bregma: CeA (coordinates in mm: anteroposterior, −1.2; mediolateral, ±2.95; dorsoventral, 4.60), dMT (anteroposterior: −1,06; mediolateral, 0; dorsoventral, 3.8 at a 16° angle relative to bregma, and the PBN (anteroposterior, −4.75; mediolateral, ±1.40; dorsoventral, 3.5). All subjects received a 10 mg/kg ketoprofen (AlliVet, St. Hialeah, FL) injection as a perioperative analgesic, and additional post-operative treatment with ketoprofen was maintained for 48 hours post-surgery.

For deep brain imaging of CeA, viral injection with AAV-expressing GCaMP7f was followed by the lowering of a sterile 25-gauge needle into the brain up to a depth of 4.30 mm from the surface to clear a path for the lens. The GRIN lens (0.6 × 7.3 mm, GLP-0673, Inscopix) was then slowly lowered to a depth of 4.45 mm from the cortical surface. The lens was secured in place with Metabond and dental cement. Mice were allowed to recover for at least two weeks following surgery before they were checked for GCaMP expression and experiments were performed at a minimum of 6 weeks following surgery.

### Behavior Paradigm

The animals were habituated to experimenter handling and the miniscope for at least 3 days prior to the test. The animals were administered with Vehicle or CP55490 (0.5 mg/kg, Cayman Chemical, catalog # 13608) through intraperitoneal (i.p.) injection at a volume of 1 ml/kg in a formulation containing ethanol, Kolliphor and saline in the ratio 1:1:18 (Sigma-Aldrich, St. Louis, MO). The injections were done 2 hours prior to the post-Veh/CP sessions. The animal was placed in the recording arena (evenly illuminated with an intensity of ∼20Lx) after the microscope was attached through an active commutator (Inscopix). The animal behavior was recorded and tracked with a top-view camera @30 FPS using ANYmaze tracking software (Stoelting Co., Wood Dale, IL) while simultaneously recording neuronal calcium activity. Each imaging session was externally triggered by TTLs delivered to the Inscopix nVue DAQ box by ANYmaze digital interface connected to ANY-maze tracking software to ensure the calcium imaging data was time-locked with the animal behavior tracking. The spontaneous neuronal activity was acquired in the animal’s home cage for 15 minutes. For predator-odor exposure sessions, the arena was a fresh home cage with bedding that had two clean 60mm glass petri-dish placed in the opposite corners of the length of the cage. A baseline session of 10 min was recorded. Following that, the predator odor, 0.25 microliter 2MT, was introduced in one of the two glass plates on a small piece of filter paper attached to it. The predator odor session was run for 20 minutes. After the vehicle session was recorded, the animals were returned to their home cage. Post-CP predator odor session was recorded 24 hours after (repeating the abovementioned procedure). For each behavior session, standard animal-pose estimation metrics for all time stamps were acquired from ANY-maze such as mobile vs. immobile, freezing, etc. For the predator odor session, the arena was divided into thirds lengthwise (fig. 3A) and the near-odor and far zone occupancy was defined for each animal as the nearest and farthest zone occupied respectively.

### Calcium Signal Acquisition and Analysis

All in-vivo calcium signals from GCaMP7f expressing neurons in CeA of freely behaving mice were acquired using the nVue miniature microscope (nVue, Inscopix) through the Inscopix acquisition software (IDAS) at 10 frames per second. LED (475nm) power, imaging gain, and the focal plane were determined for each animal on the first imaging session and kept constant across all subsequent sessions. The recorded calcium imaging data was first processed using the Inscopix data processing software (v 1.9.5 Inscopix). Briefly, data was preprocessed, bandpass filtered, and corrected for movement artifacts. Calcium traces from individual neurons were extracted using constrained non-negative matrix factorization for the endoscopy algorithm (CNMFe)^54^ and false positives were manually excluded from further analysis. Longitudinal registration across sessions was performed in IDPS with a minimum correlation of 0.5 as threshold. In a subset of mice, GCaMP was expressed under synapsin promoter in SOMAi14 mice, and SOM cells were identified by overlaying the extracted cells with the spatial map obtained from the red channel. For home cage spontaneous calcium activity analysis, the raw dF/F traces were deconvolved using OASIS^55^ followed by the calculation of event frequency, mean event amplitude, and area under the curve (AUC) using codes written in MATLAB.

The linear dependence between the activity of two neurons was quantified by computing their Pearson correlation coefficient (CC) on the 10-minute time series of denoised Ca^2+^ recordings by using the MATLAB function *corcoeff*. Interactions between simultaneously recorded populations of SOM neurons were examined using network theoretic metrics by using custom-written codes in MATLAB and the Brain Connectivity Toolbox (BCT)^56^. We first build an adjacency matrix *(A)* with the active neurons denoting ‘nodes’ in the network. The weight of each entry of the matrix, *Aij*, was the CC between Ca^2^ traces of neurons *i* and *j*. The sign of CC values was preserved. We set a threshold cutoff of |CC| > 0.3 to define whether a ‘link’ or ‘connection’ exists between a pair of neurons as has been used previously as a reasonable estimate of connectivity. Separate adjacency matrices were built for positive and negative connections. Connection density was defined as the fraction of present connections to possible connections. The degree (*k*) of a node was the number of links it had with other nodes in the network and the degree distribution (*p_k_*) provided the probability that a randomly selected node in the network had degree *k*. The strength (*s*) of a node was the sum of the weights of links connected to the node and its probability density was *p_s._* The sparsity index was calculated as the fraction of non-zero entries in A. Centrality was measured using the ‘eigenvector’ centrality function which uses the eigenvector corresponding to the largest eigenvalue of A. Circular maps of networks, in which the nodes were organized in the circumference and connections drawn within the circle, were created using the MATLAB function *circularGraph*.

We sought to identify whether these networks could be partitioned into nonoverlapping groups or modules by implementing the Louvain community detection algorithm^56,57^. We implement this optimization algorithm by combining positive and negative connections in one matrix and by utilizing symmetric treatment of weights. This maximizes the number of within-group positive links and minimizes within-group negative links, while simultaneously minimizing the number of between-group positive links and placing nodes that have negative links with the same group of nodes in one module. We parallelly identified highly connected nodes in a network by defining them to be the nodes whose total degree is +1 standard deviation more than the mean of all nodes in the network and called them hubs. We then used the module assignments to calculate the participation coefficient (PC)^58,59^ which measures how ‘well-distributed’ the links of a node are among different modules and is formally defined as follows:

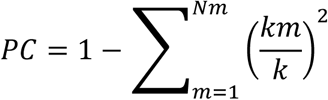

Where *Nm* is the number of modules, *k* is the degree of the node and *km* is the number of connections from the node to module *m*. PC is close to 1 if a node is connected uniformly among all the modules, and 0 if all its connections are within its own module. Hubs and their participation coefficient were calculated separately for positive and negative connectivity matrices. Modularity and PC were computed using BCT in MATLAB.

For predator-odor sessions, neuronal activity during behavior transitions such as odor approach, flee, and immobility/freezing were quantified in MATLAB. Single neuron traces were aligned to each behavior onset bout (zone approach, flee, freezing onset respectively), isolated (from 2s before onset to 5s after), and averaged across all bouts of each behavior class. The averaged traces were then Z-transformed using the signal from 2 to 1.5s before the behavior onset as a baseline. The Traces exceeding a Z-score value of 2.58 or -2.58 (p = 0.01, two-tailed) for any 2 consecutive frames in a window of 2s following behavior onset were either considered (+) Responsive, (-) Responsive respectively or otherwise unresponsive. The proportion of cells satisfying each of the three criteria was then formally compared between vehicle and CP55490 sessions using a Chi-squared test of proportions. The AUC for the activity of neurons in each group during pre and post-behavior onset was also quantified and compared. The behavior selectivity and relative overlap of neuronal activity were further analyzed using Euler-Venn diagrams (created using a code written in Python).

To test if the treatment used (i.e. Vehicle or CP55490) during predator odor (2MT) exposure sessions could be decoded from peri-event instantaneous CeA neuronal activity associated with behavioral transitions, we trained and tested linear support vector machine (SVM) classifiers with a Gaussian kernel in MATLAB. We independently trained and tested classifiers for each 0.1 s separately in the following manner. Data matrices corresponding to each session type were randomly split into training and test datasets (3:2 ratio). From these, training and testing datasets we randomly selected 100 values this was done iteratively 50 times to create 100 x 50 matrices for each session type and training and testing datasets. The resulting training matrices were vertically concatenated to create a 100 x 100 training matrix which was fed into the SVM together with the session identity of each row. Classifiers were validated using five-fold cross-validation. A similar procedure was performed to create a testing dataset matrix and fed onto the SVM classifier and accuracy was quantified as the proportion of correct classifications. This procedure was repeated 10 times to determine average classifier performance accuracy and variability. The decoder accuracy was compared to decoder accuracy obtained from training and testing temporally permuted time series datasets in a similar manner to that described above. The permuted data set was obtained by circularly permuting the time series from each neuron randomly for 100 iterations.

Principal component analysis (PCA) was conducted using available MATLAB functions to identify population dynamics^60^ for each behavior transition as described above. Bout averaged Z-scored traces for each behavior transition were concatenated together for vehicle and CP55490 sessions. Since the number of detected neurons in the CP55490 sessions using CNMFe was consistently higher than those of vehicle sessions in all animals, traces for the CP session were randomly selected to create a matrix that had the same number of traces as the one for the vehicle session. PCA was applied to the concatenated matrix to reduce dimensionality for further assessment. For visualization, the first 3 PC dimensions for each group and behavior type were plotted (Fig 2n, Fig. S6). We then compared the distance of freeze onset and Flee onset to Approach onset population trajectory in PC space using Mahalanobis distance (MD) between approach-flee and approach-freeze traces respectively for each session using a resampling and iterative method described below. We additionally compared the MD between vehicle and CP sessions for all possible behavior-pairs (i.e approach-flee, flee-freeze, approach-freeze). To calculate MD, we randomly sampled 50 % of the neurons from each session (100 repetitions) and used the number of PCs that accounted for 90% of the cumulative variance in the dataset. The resampling method was used to factor in the differences in the number of cells detected for each session type. The temporal profile of MD between each behavior duo was then formally compared for vehicle and CP sessions.

Agglomerative hierarchical clustering was applied to cluster neurons based on their activity profile during the three behavior transitions^28^. Briefly, Z-scored traces from both vehicle and CP sessions were pooled together. The time dimension of the traces was transformed using PCA and then clustered using correlation as the distance metric. The number of optimal clusters, 8 in our case, was determined based on the optimal cluster number, which is associated with the largest gap value of the Calinski–Harabasz Index^61^. The traces within each cluster were then split into their session ID (i.e. vehicle or CP). The relative member proportion and AUC of each cluster were calculated and formally compared.

We calculated area or mobility bias in single neuron responses during predator-odor exposure sessions. The epochs that defined the animal’s presence near/far away from the odor along with the mobile/immobile states were acquired from ANYmaze software. Single neuron traces for the entire session were Z-normalized. A neuron was considered active if the trace surpassed +1.96 Z scores. Activation probability was then calculated for each area occupancy/ mobility scenario using the following formula:

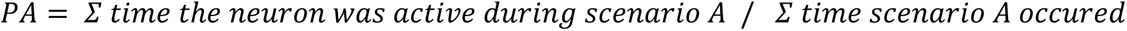

To calculate the bias, two indices were calculated as follows:

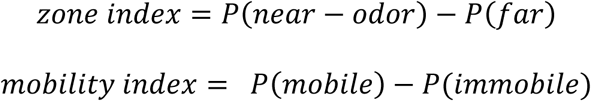

A neuron was considered selective if its index value was either > +2 SD or < -2 SD of the shuffled index (using *randperm* function in MATLAB to produce a temporal scrambled set of behavior/zone occupancy markers) for the same session. The proportion of neurons that were selective for a particular scenario was then formally compared across vehicle and CP sessions.

### Histology

To confirm location of GCaMP expression and lens location, mice were transcardially perfused with 1× PBS followed by 4% paraformaldehyde solution. Brains were fixed overnight at 4 °C, and then transferred to 30% sucrose solution for 48 h. 50 μm coronal sections of the brains were obtained using a vibratome (Leica VT1000 S) and mounted on glass slides with VectaShield H-1200 DAPI mounting medium (Vector Laboratories, Burlingame, CA). Images were obtained using an epifluorescent microscope (Keyence BZ-X800) with a 10× objective.

### Ex vivo Electrophysiology

Coronal brain sections were collected at 250 μm using standard procedures. Mice were anesthetized with isoflurane and transcardially perfused in an ice-cold/oxygenated (95% v/v O_2_, 5% v/v CO_2_) cutting solution consisting of (in mM): 93 N-methyl-D-glucamine; 2.5 KCl, 20 HEPES; 10 MgSO_4_. 7H_2_O; 1.2 NaH_2_PO_4;_ 30 NaHCO_3_; 0.5 CaCl_2_. 2H_2_O; 25 glucose; 3 Na+ pyruvate; 5 Na+ ascorbate; and 5 N-acetylcysteine. The brain was subsequently dissected, hemisected, and sectioned using a vibrating Leica VT1000 S microtome (Leica Biosystems). The brain slices were then transferred to an oxygenated 34 °C chamber filled with the same cutting solution for a 10-minute recovery period. Slices were then transferred to a holding chamber containing a buffered solution consisting of (in mM): 92 NaCl; 2.5 KCl; 20 HEPES; and 2 MgSO_4_. 7H_2_O; 1.2 NaH_2_PO_4_; 30 NaHCO_3_; 2 CaCl_2_. 2H_2_O; 25 Glucose; 3 Na+ pyruvate; 5 Na+ ascorbate; and 5 N-acetylcysteine. They were allowed to recover for ≥30 minutes. For recording, slices were placed into a perfusion chamber where they were constantly exposed to oxygenated artificial cerebrospinal fluid (31–33 °C) consisting of (in mM): 113 NaCl; 2.5 KCl; 1.2 MgSO_4_. 7H_2_O; 2.5 CaCl_2_. 2H_2_O; 1 NaH2PO_4_; 26 NaHCO_3_; 20 Glucose; 3 Na+ pyruvate; and 1 Na+ ascorbate at a flow rate of 2–3 ml min^−1^.

Drug concentrations for electrophysiology experiments were (in μM): 10 Rimonabant (Cayman Chemical, catalog #9000484), 5 CP55940 (Cayman Chemical, catalog # 13608), 1 Gabazine (Sigma-Aldrich, catalog # S106), 50 Picrotoxin (Sigma-Aldrich, catalog #528105). Drug incubations were performed for 30-60 min at room temperature prior to recordings. Drug wash-on experiments were carried out after 5 minutes of stable baseline recording. Stock solutions of CP and RIM were prepared in DMSO, and final recording solutions contained 0.05% w/v Bovine Serum Albumin (BSA, Sigma-Aldrich, St. Louis, MO, USA).

Cells were visually identified from Ai14 reporter lines or virally injected animals under illumination from a series 120Q X-Cite lamp at ×40 magnification using an immersion objective. Neurons were voltage-clamped in whole-cell configuration using borosilicate glass pipettes (3– 6 MΩ) filled with intracellular solution containing (in mM): 120 CsOH, 120 D-gluconic acid, 2.8 NaCl, 20 HEPES, 5 TEA-Cl, QX314-Cl 5 mM, 2.5 Mg-ATP, 0.25 Na-GTP (pH 7.30–7.35, Osmolarity: 285-290 mOsm). Current clamp recordings were performed with a K^+^-gluconate internal solution (in mM): 125 K+ -gluconate, 4 NaCl, 10 HEPES, 4 MgATP, 0.3 Na-GTP, and 10 Na-phosphocreatine (pH 7.30–7.35, Osmolarity: 280-290 mOsm). Following break-in to each cell, at least 3 min of time elapsed before the start of the experiments to allow for internal solution exchange and stabilization of membrane properties. Bridge balance was applied to correct series resistance online in the current clamp mode.

All data were acquired using a MultiClamp 700B amplifier equipped with Digidata-1550A A/D converter (Molecular Devices); data were acquired at 10 kHz and low-pass filtered at 1 kHz and analyzed with the Axon pCLAMP 10 electrophysiology data acquisition and analysis software version 10.7 (Molecular Devices). The liquid junction potential was not compensated. Access resistance was monitored throughout the experiments, and neurons in which the series resistance exceeded 30 MΩ or changed by 20% or more were excluded from the statistics.

mEPSC and mIPSC recordings: Continuous current traces of 5-min duration at -60 mV were recorded in the presence of Tetrodotoxin citrate (TTX, 1μM, Tocris Bioscience) to obtain mEPSCs, recorded at least 3 min after achieving whole-cell configuration. Cells were then held at +15 mV and after the cell holding current stabilized, mIPSCs were obtained as a 5 min continuous recording.

### Ex vivo optogenetics

Mice recovered for 5-7 weeks following virus injection. Optically evoked responses were recorded from CeA SOM neurons ipsilateral to the hemisphere where injections were performed. Photostimulation was achieved through a 40x objective at 0.03-0.05 Hz using a 455 nm LED system (Mightex, BLS-Series, Toronto, Canada), which was controlled via the analog output of the Digidata. Stimulation duration ranged between 0.5-2 milliseconds and LED power between 0.6-1.5 mW. EPSCs and IPSCs were isolated by recoding at holding potentials of −60 mV and +15 mV, which were the empirically derived reversal potentials for GABA and glutamatergic input respectively. The effect of CP55940 wash-on on current amplitude were calculated as a percent of, and compared to, a (5-minute) pre-stimulation baseline. PPR was also obtained in these recordings with an inter-stimulus interval of 50 milliseconds. CP55940 sensitivity of local inhibition (Fig 5) was measured in a similar manner as above while selectively expressing ChR2 in SOM^+^ (Cre-on) or SOM^-^ neurons (using a Cre-off strategy^53^) and recording IPSCs from SOM^+^ cells at +10 mV (reversal potential for ChR2), to minimize ChR2 driven inward currents.

We next tested the effect of local inhibitory input activation on the spiking of SOM neurons and its sensitivity to CP, again by expressing ChR2 in SOM^-^ neurons as mentioned above. This induced short latency IPSPs in SOM^+^ cells upon optical stimulation, recorded while injecting current to hold the cell at -55 to -57 mV. Spiking in the SOM^+^ cells was achieved by depolarizing the cell body through the injection of square pulses. The pulses were composed of a sequence of five short (10 ms) current steps instead of a longer single step to achieve the dual purpose of minimizing effects due to spike adaptation or channel desensitization; and organizing the pulses at different inter-pulse intervals to mimic different frequencies of input stimulation (5, 10 and 20 Hz). The amount of current injected (range: 100 to 300 pA) was determined on a cell-to-cell basis and it was the minimum tested value required to reliably induce firing at the first 10 ms pulse. The recordings alternated between trials where 5 ms blue light were delivered 2 ms before start of the current pulse and those which only had current injections, and the light flashes were not triggered. Each AP is identically weighted in its contribution to the AP probability, averaged over 10 trials. For instance, a cell’s AP probability will be 1 if fires an AP in all 5 pulses in all 10 trials. Similarly, time-varying sinusoidal current pulses with peak amplitude between 100 to 300 pA and frequency of 3.5-5 Hz were injected, which constrained the AP occurrence to a narrow time window at only the peaks of the wave^62^. The timing of optical activation of SOM-population was performed in reference to the positive peak of the third oscillation cycle while recording from SOM^+^ cells. This is described as Δt and negative values indicate light stimulation prior to the peak.

### Ex vivo Ca^2+^ imaging

GCaMP7f fluorescence of CeA SOM in brain slices was monitored using a camera equipped with a CMOS sensor (DS-Qi2, Nikon, Melville, NY) mounted on a Nikon Eclipse FN-S2N microscope. Ex vivo brain slice preparation, recovery, and holding for Ca^2+^ imaging were performed as detailed above for electrophysiological experiments, with an additional step of reintroduction of Na+ during recovery for improved viability^63^. Photostimulation was performed using a Mightex 455 nm LED system (BLS-Series) (Mightex, Toronto, Ontario, Canada). ROI of 512 X 512 pixels was imaged under constant illumination for 5 mins at 10 Hz through a 40× water-immersion objective using NIS-Elements D software. Bleach correction of the recordings was performed using the ‘exponential fit’ method in Fiji^64,65^. Subsequent analysis was performed Inscopix Data Processing Software pipeline (IDPS), using the PCA-ICA module to detect Ca^2+^ events.

### Statistical analysis

Statistical analyses and generation of graphs were performed in Prism 10 (GraphPad Software) and MATLAB (R2023b). For the analysis of two groups, an unpaired or paired Student’s t-test was used if data was normally distributed, otherwise non-parametric Mann-Whitney U test was used for non-normal data. Normality was tested using the D’Agostino & Pearson Anderson-Darling or Shapiro-Wilk test. For analysis of three or more groups across a single independent variable, a one-way ANOVA was used with a Holm-Sidak posthoc multiple comparisons test between groups as noted in the figure legends. For analysis between two or more groups across two or more independent variables, a two-way ANOVA was used with a Tukey’s or Holm-Sidak posthoc multiple comparisons test between groups as noted in the figure legends. For the comparison of distributions, the two Kolmogorov-Smirnov tests were used (K-S test). Proportions were compared using the Chi-square test. The threshold for statistical significance was set at P<0.05 in all datasets. No statistical methods were used to predetermine sample sizes, but our sample sizes are comparable to previous publications.

## ACKNOWLEDGEMENTS

These studies were supported by NIH grants MH100785 (S.P.) and K08 MH126166 (L.E.R-V.) and NARSAD Young Investigator Awards (F.Y., L.E.R-V., and S.N.). We acknowledge technical assistance provided by Meredith Cone, Danyal Zaidi, Isaac Kandil, and Pheobe Cao.

## AUTHOR CONTRIBUTIONS

S.P. and F.Y. conceived the work, designed experiments, and interpreted data. F.Y. and S.N. designed, executed, and analyzed experimental data. L.E.R-V. analyzed data. All authors contributed to manuscript preparation.

