## Supplemental File for "Cannabinoid Modulation of Central Amygdala Population Dynamics During Threat Investigation"

### SUPPLEMENTARY FIGURES

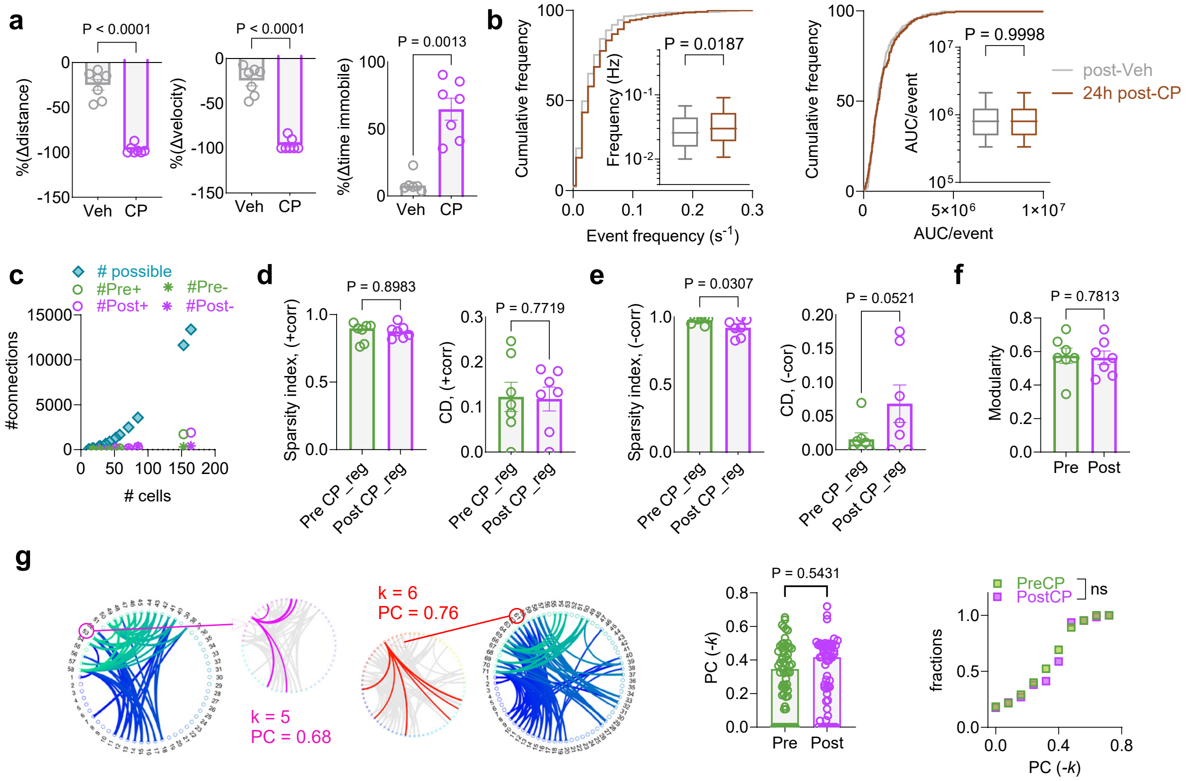

**Figure S1: a)** CP administration reduces total distance travelled and mean velocity and increases duration of immobility in comparison to Veh (distance: t=12.42, df=6; velocity: t=11.82, df=6; immobility: t=5.660, df=6; Paired t-test; N=7 mice). **b)** Cumulative frequency histograms and box plots of event frequency and AUC/event showing comparison of post-Veh with 24 hrs post-CP (P value via Mann Whitney test, $U_{freq}$ = 38469, $U_{AUC}$ = 36181, N = 3 mice). **c)** XY scatterplot with number of cells observed on the X-axis, the number of connections possible, and the number of connections observed (pre and post, positive and negative) on Y-axis for all 7 mice. **d)** No significant difference in sparsity index or connection density of positive connections in the cells registered across both pre- and post-CP sessions (P value via paired t-test, SI: t=0.1333, df=6; CD: t=0.3033, df=6). **e)** Reduction in sparsity index of negative connections and trend towards enhanced connection density in the cells registered across both pre- and post-CP sessions (P value via paired t-test, SI: t=2.812, df=6; CD t=2.416, df=6). **f)** No significant change in community structure defined via modularity pre- and post-CP (P value via paired t-test, t=0.2904, df=6). **g)** Circular graph representation of negative connections of the same network as presented in Fig.1o, with a negative hub being highlighted in pre- and post-CP sessions. No change in participation coefficients (PC) of negative hubs was observed (P value via unpaired t-test, t=0.6098, df=126).

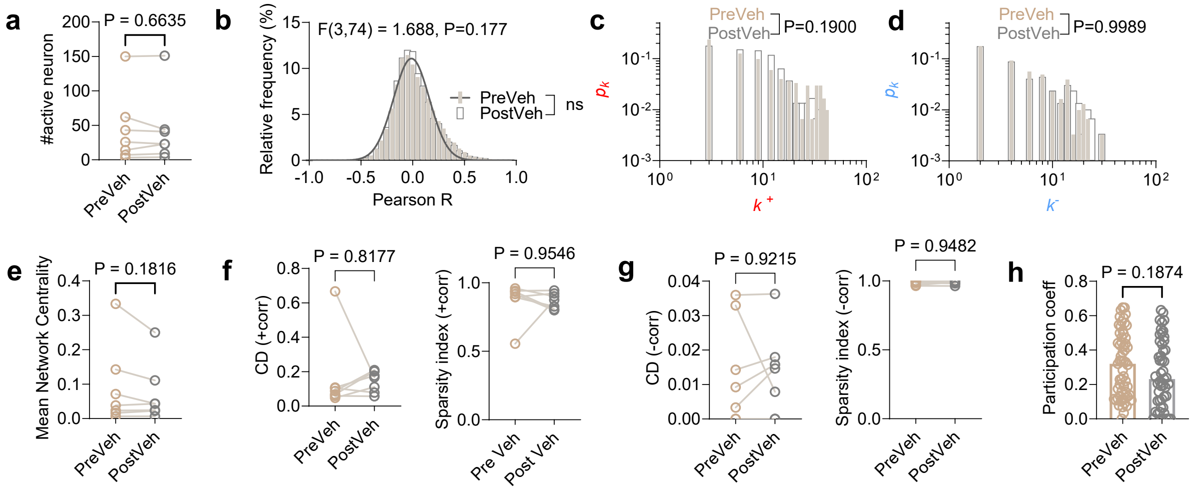


**Figure S2: a)** Number of active cells in pre-Veh and post-Veh sessions (Paired t-test, t=0.4574, df=6 N=7 mice). **b)** Distribution of Pearson's correlation coefficients for all simultaneously recorded pairs of neurons along with gaussian fit (goodness of fit, $R^2$=0.97) with least square regression (F and P value via fit comparison with Extra sum-of-square F test). **c)** Degree distributions of positive connections pre- and post-Veh (P value via KS test, D=0.0886). **d)** Degree distributions of negative connections pre- and post-Veh (P value via KS test, D=0.02879). **e)** Mean network centrality is not affected by Veh treatment. (P value via paired t-test, t=1.511, df=6). **f)** Connection density and Sparsity index of networks constructed from positive connections (P value via paired t test, CD: t=0.2409, df=6, SI: t=0.059, df=6). **g)** Same as (f), for negative correlations. (P value via paired t tests, CD: t=0.103, df=6; SI: t=0.068, df=6). **h)** PC values of hubs in networks are unchanged pre- and post-Veh (P value via unpaired t-test, t=1.327, df=106).

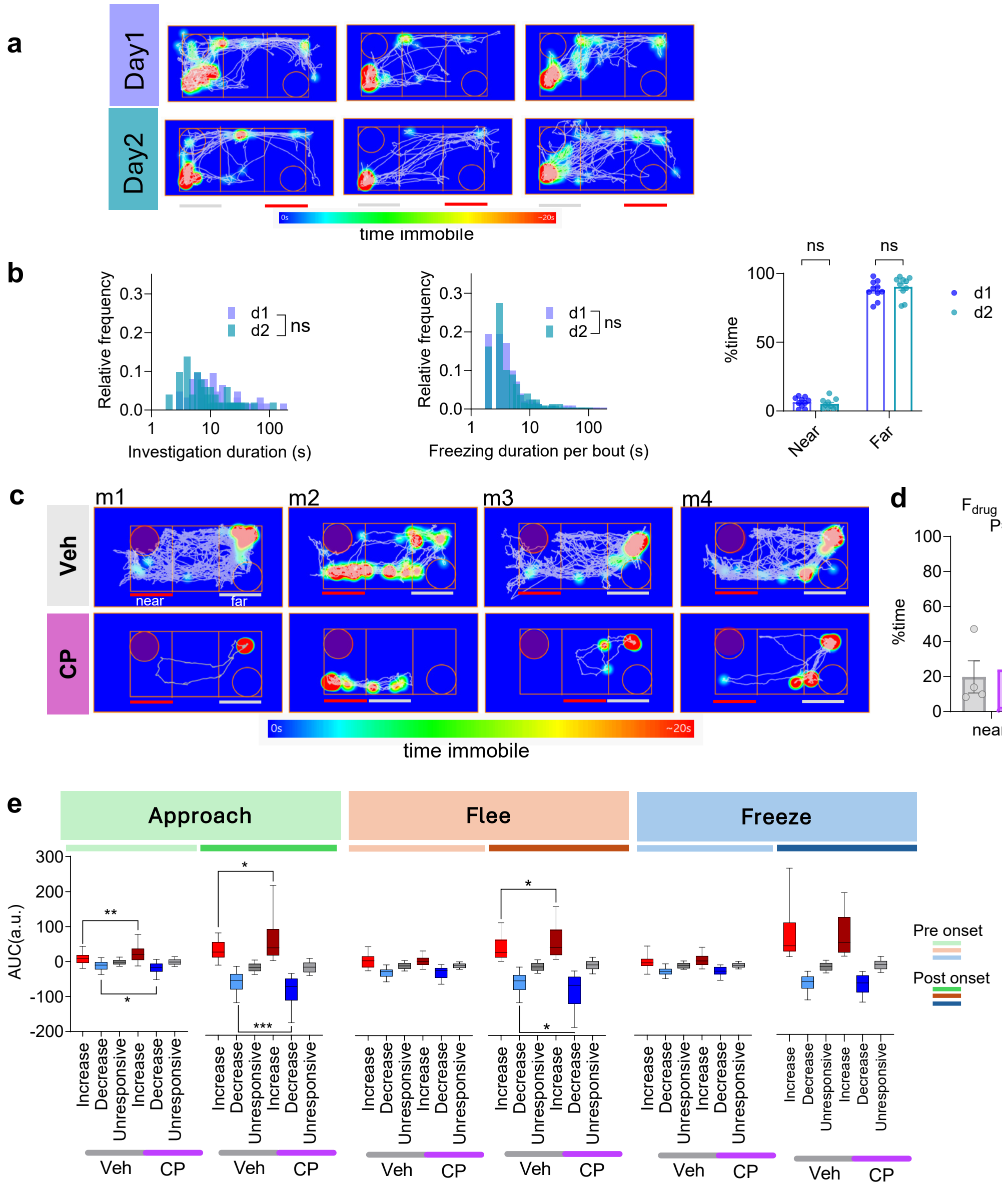


**Figure S3: a)** Locomotion trajectory (grey lines) and immobility (heatmap overlay) of animals subjected to 2MT exposure for two consecutive days. The grey and red lines at the bottom refers to the far zone and near-odor zone respectively. **b)** Distribution of time spent investigating the predator odor zone for all approach bouts (left, KS test, D=0.2502), freezing bout length (center, KS test, D= 0.05485) and total % time spent freezing (right, 2-Way RM ANOVA, $F_{d1/d2}$ (1, 9) = 0.4724, P=0.5092, Tukey's posy hoc multiple comparison) during both 2MT sessions. **c)** Locomotion trajectory (grey lines) and immobility (heatmap overlay) of 4 tethered mice subjected to 2MT exposure post vehicle (top) and CP (bottom) administration. The grey and red lines at the bottom refers to the far zone and near-odor zone respectively specified for each animal based on its individual trajectory path. For m3, for example, near zone was the intermediale third of the chamber as it never ventured into the third of the chamber closest to the odor. **d)** Percent time spent in each zone during 2MT exposure for vehicle (grey) and CP (purple) session; F and P values from 2-way RM ANOVA. **e)** Area under the curve for each group of neurons that are either responsive (increase/decrease activity) or unresponsive for each behavior onset class. The AUCs are segregated into pre-onset and post-onset. Descriptions of statistically significant comparisons between Veh and CP via Mann-Whitney test: Approach: $U_{pre\_Increase}$=1909, **P=0.0026, $U_{pre\_decrease}$=5601, *P=0.0191, $U_{post\_Increase}$= 2085, *P= 0.0197, $U_{post\_decrease}$= 4940, ***P= 0.0003. Flee: $U_{post\_Increase}$= 2978, *P= 0.0187, $U_{post\_decrease}$= 2345, *P= 0.0229.

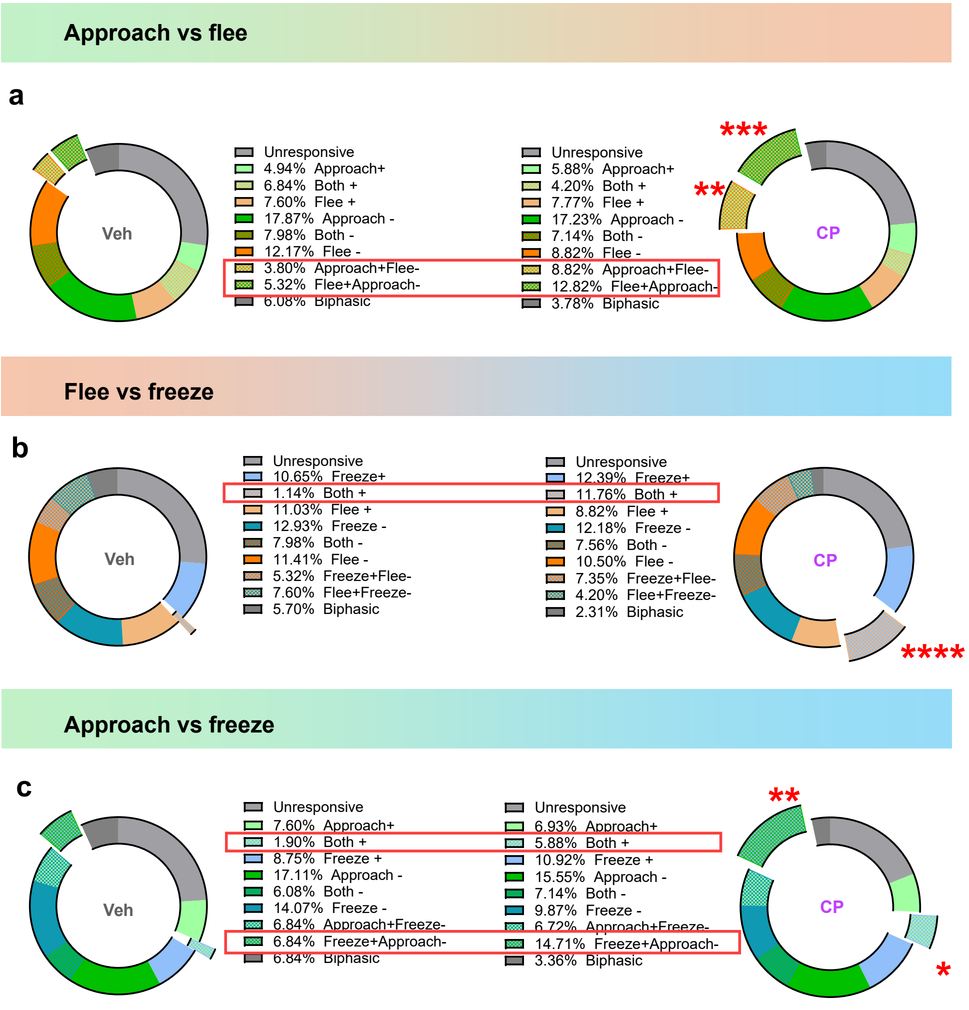


**Figure S4:** Comparision of cell proportions responsive (Z-score >±2 SD from baseline) to pairs of behaviours during 2MT exposure in Veh and CP-treated mice using Chi square test for proportionality. The statistically significant groups are highlighted. **a)** Approach+Flee– ($\chi$ = 6.53, **P = 0.01), Flee+Approach– ($\chi$ = 10.43, ***P = 0.001). **b)** Both+, i.e. positive for both flee and freeze ($\chi$ = 26.03 ****P=3.37e-07). **c)** Freeze+Approach– ($\chi$ = 9.98, **P = 0.0016), Both +: ($\chi$ = 6.29, *P = 0.012).

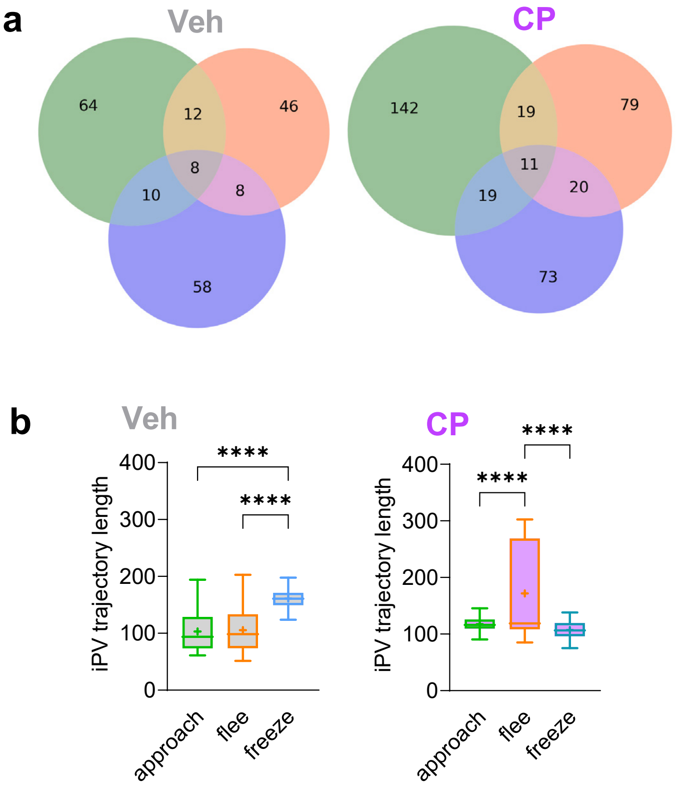


**Figure S5: a)** Venn diagram to show overlap in the number of cells that are negatively responsive for behavioral onset of approach, flee and freeze [Significantly different group between Veh and CP, '(-) Freeze selective neurons': Freeze - (Approach ∩ Freeze) - (Flee ∩ Freeze) - (Approach ∩ Flee ∩ Freeze): Veh=58, CP=73, $\chi$ = 4.7996, P = 0.0285, Chi square test]. **b)** Length of neural population trajectory associated with the described behavioral bouts in Veh animals (Ordinary one-way ANOVA, F(2,297)=117.7, P<0.0001, p values on graph from Tukey's multiple comparison) and CP (Ordinary one-way ANOVA, F(2,297)=53.98, P<0.0001, p values on graph from Tukey's multiple comparison)

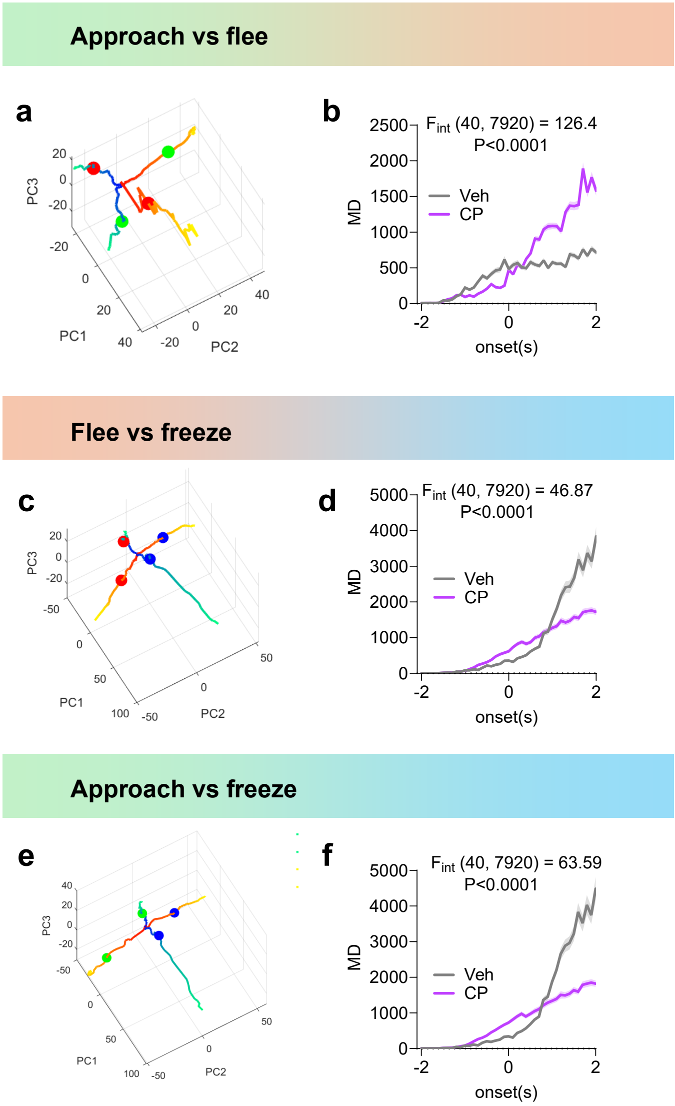


**Figure S6:** Visualization of neural trajectories in the same state space for pairs of behaviors during 2MT exposure in vehicle or CP-treated mice via pooling all neurons for dimensionality reduction and then separating them into Veh and CP for comparison. **a)** Neural trajectories for approach or flee, onset denoted by green and red circles respectively. Blue-green trajectories: Vehicle; Red-Yellow trajectories: CP. **b)** Mahalanobis distance (MD) between approach and flee trajectories for vehicle (grey) and CP (purple) treated animals. **c)** Neural trajectories for flee or freeze, onset denoted by red and blue circles respectively. Blue-green trajectories: Vehicle; Red-Yellow trajectories: CP. **d)** MD between flee and freeze trajectories for vehicle (grey) and CP (purple) treated animals. **e)** Neural trajectories for approach or freeze, onset denoted by green and blue circles respectively. Blue-green trajectories: Vehicle; Red-Yellow trajectories: CP. **f)** MD between approach and freeze trajectories for vehicle (grey) and CP (purple) treated animals. F and P values in (b,d,f) via 2-way RM ANOVA.

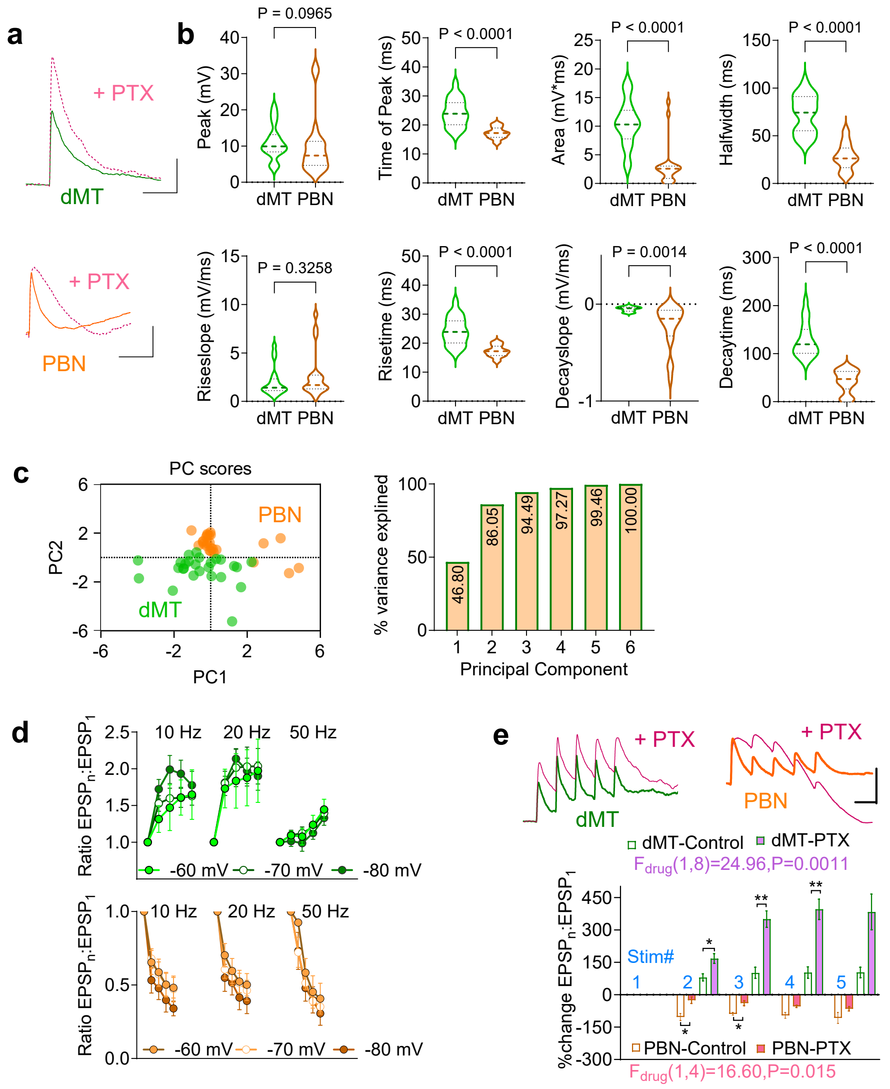


**Figure S7**: **a)** Example traces of EPSPs evoked by stimulation of dMT (top) or PBN (bottom) inputs. Dotted traces are EPSPs recorded from the same cell after $GABA_A$ receptor blockade by picrotoxin. (Scale bar: 5 mV, 100 ms). **b)** Violin plots of comparisons between EPSP features evoked by dMT or PBN inputs. Unpaired t-test for comparison of halfwidth (t=8.145, df=38), decay slope (t=3.439, df=38), decay time (t=8.877, df=38), rise slope (t=0.9955, df=38) and rise time (t=6.056, df=38). Mann-Whitney test for Peak (U= 138), Area (U=31) and time of peak (U=31). **c)** PC scores calculated from parameters described in b show each input occupies a distinct PC space. First 3 PCs explained 94% of the cumulative variance. **d)** Summation of synaptically driven EPSP at different stimulation frequencies (10, 20 and 50 Hz) show facilitation for dMT (top) and depression for PBN (bottom). **e)** Example traces (top) showing the effect of blocking synaptic inhibition on EPSP summation at 20 Hz stimulation for dMT and PBN (Scale bar: 5 mV, 50 ms). Quantification (bottom) of enhanced degree of summation for dMT and reduced degree of depression for PBN input after picrotoxin (PTX) (F and P values via 2-way RM ANOVA, Tukey's post hoc, *p<0.05, **p<0.01).

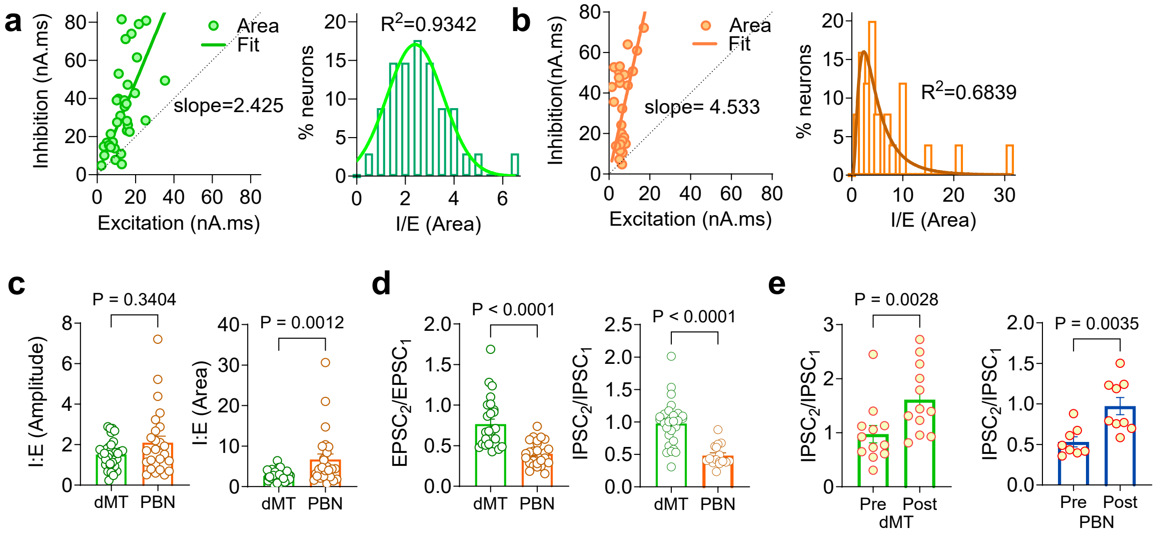


**Figure S8: a)** XY scatter plot of EPSC and IPSC area under the curve, with the fit computed via linear regression. Summary of IE ratio, which was best fit by a gaussian model, $R^2$ for goodness of fit. **b)** Same as (a) for PBN input stimulation, frequency distribution was best fit by lognormal model. **c)** Inhibition-excitation (I:E) ratio at dMT-CeA and PBN-CeA synapses, analyzed using peak amplitude (left) and area (right) (Mann-Whitney test, $U_{amp}$=362, $U_{area}$=218). **d)** Comparison of basal PPR at the excitatory inputs elicited by dMT or PBN stimulation (left, Mann-Whitney test, U=76). Difference in basal PPR of inhibitory currents (right, Mann-Whitney test, U=41). **e)** Significant enhancement of PPR calculated for inhibitory currents pre- and post-CP wash at dMT input (left, Paired t test, t=3.828, df=11) and PBN input (right, Unpaired t test, t=3.460, df=15).
